# The importance of charge in perturbing the aromatic glue stabilizing the protein-protein interface of homodimeric tRNA-guanine transglycosylase

**DOI:** 10.1101/2020.09.01.277731

**Authors:** Andreas Nguyen, Dzung Nguyen, Tran Xuan Phong Nguyen, Maurice Sebastiani, Stefanie Dörr, Oscar Hernandez-Alba, François Debaene, Sarah Cianférani, Andreas Heine, Gerhard Klebe, Klaus Reuter

## Abstract

Bacterial tRNA-guanine transglycosylase (Tgt) is involved in the biosynthesis of the modified tRNA nucleoside queuosine present in the anticodon wobble position of tRNAs specific for aspartate, asparagine, histidine and tyrosine. Inactivation of the *tgt* gene leads to decreased pathogenicity of *Shigella* bacteria. Therefore, Tgt constitutes a putative target for Shigellosis drug therapy. Since only active as homodimer, interference with dimer-interface formation may, in addition to active-site inhibition, provide further means to disable this protein. A cluster of four aromatic residues seems important to stabilize the homodimer. We mutated residues of this aromatic cluster and analyzed each exchange with respect to dimer and thermal stability or enzyme activity applying native mass spectrometry, thermal shift assay, enzyme kinetics, and X-ray crystallography. Our structural studies indicate strong influence of pH on homodimer stability. Obviously, protonation of a histidine within the aromatic cluster promotes the collapse of an essential structural motif within the dimer interface at slightly acidic pH.

**TOC Graphic:** For table of contents use only.

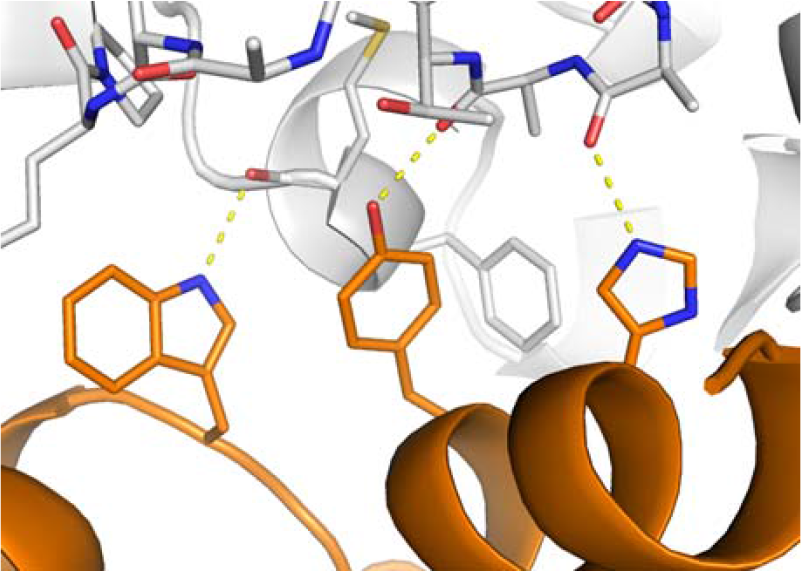

## Introduction

In the majority of bacteria and eukaryotes, the anticodon wobble position (34) of tRNA^Asp^, tRNA^Asn^, tRNA^Tyr^ and tRNA^His^ is occupied by the highly modified nucleobase queuine, which is involved in the fine-tuning of translational accuracy and speed.^1,2^ Only bacteria possess the enzymes required for the complex biosynthesis of queuine. It starts outside the tRNA with GTP being converted to 7- (aminomethyl)-7-deazaguanine by five enzyme-catalyzed reactions.^3–7^ Accompanied by the excision of the genetically encoded guanine, this precursor, which is also referred to as preQ_1_ base, is then inserted into position 34 of the above-named tRNAs in a reaction catalyzed by the enzyme tRNA-guanine transglycosylase (Tgt; EC 2.4.2.29).^8^ Thereupon, at the tRNA level, two further enzymes are required to accomplish the final synthesis of the queuine nucleoside, queuosine.^9–13^

In *Shigella* spp., the causative agents of bacillary dysentery, inactivation of the *tgt* gene compromises the translation of *virF* mRNA encoding a transcriptional activator of numerous essential virulence genes.^14^ The resultant attenuation of the pathogenicity of this organism provided the opportunity to use bacterial Tgt as an object for the rational design of compounds with therapeutic potential against Shigellosis. Investigated by high-resolution crystal structures of the *Zymomonas mobilis* orthologue,^15^ bacterial Tgt has, therefore, probably become the most intensively studied tRNA-modifying enzyme to date. Its functional unit consists of a stable homodimer with the protomer adopting a (βα)_8_ barrel fold. This, however, deviates from the classical motif by an N- and a C-terminal extension as well as by several insertions. The most prominent of these insertions is a Zn^2+^-coordinating subdomain between β-strand 8 and α-helix 8 of the barrel motif, which is involved in the organization of the homodimer interface. Xi et al.^16^ succeeded in crystallizing *Z. mobilis* Tgt in complex with a 20-meric RNA oligonucleotide, which mimics the anticodon stem-loop of *Escherichia coli* tRNA^Tyr^. The authors determined two crystal structures of this complex, which reveal that the enzyme is able to bind only one RNA molecule at a time. Although each subunit contains an entire functional active center, both substrate-binding pockets are located on the same side of the homodimer, inevitably leading to a steric clash of two simultaneously bound tRNA substrates. Obviously, in the active complex, one monomer performs catalysis, while the second one is required for positioning the tRNA substrate in correct orientation.

In previous studies, we came across Tgt active-site inhibitors which turned out to destabilize the subunit interface of the Tgt homodimer. These inhibitors are based on a *lin*-benzoguanine scaffold, which occupies the substrate base binding pocket of the enzyme. Furthermore, each of these compounds is endowed with an extended 4-substituent, which pokes into the nearby protein/protein interface and, thus, takes impact on its geometry.^17,18^ Notably, a small subset of these “spiking ligands” is able to induce a circa 130° rotation of one subunit of the homodimer against the second one. This rearrangement leads to an unproductive quaternary structure of the enzyme, referred to as “twisted dimer”. Although there is no evidence yet if this unconventional dimer form is of physiological relevance, its stabilization may offer a promising strategy for the design of Tgt inhibitors since, for steric reasons, it is unable to bind a tRNA substrate.^19,20^

The finding that small ligands may interfere with Tgt dimer formation had led us to investigate the interface architecture of the Tgt dimer. A detailed analysis of the homodimer protein/protein interface on the basis of the *Z. mobilis* Tgt apo-structure (pdb-code: 1pud) revealed a contact area of more than 1600 Å^2^ containing 43 interface residues per subunit. In addition to hydrophobic interactions accounting for more than 60% of the interface, these residues form ten salt bridges and eight hydrogen bonds to residues of the second subunit. Comparison of the inter-subunit interactions seen in 1pud with those observed in another 43 crystal structures of *Z. mobilis* Tgt showed that most of them are present in all of these structures, but some are not, which indicates a certain plasticity of the interface. As revealed by molecular dynamics simulations and further in silico-analyses, a cluster of four aromatic residues with mutual distances of 5 to 6 Å appears to be especially important for dimer stability.^21^ Three of these amino acids, namely Trp326, Tyr330 and His333, are located in or close to α-helix E of one subunit, while the fourth one, Phe92’, resides in α-helix 2c’ of the second subunit (indicated by a “’”) (Figure 1). Due to the two-fold symmetry relating the dimer mates to each other, this cluster is present twice in the interface, as are virtually all interactions observed between both subunits.

**Figure 1:**
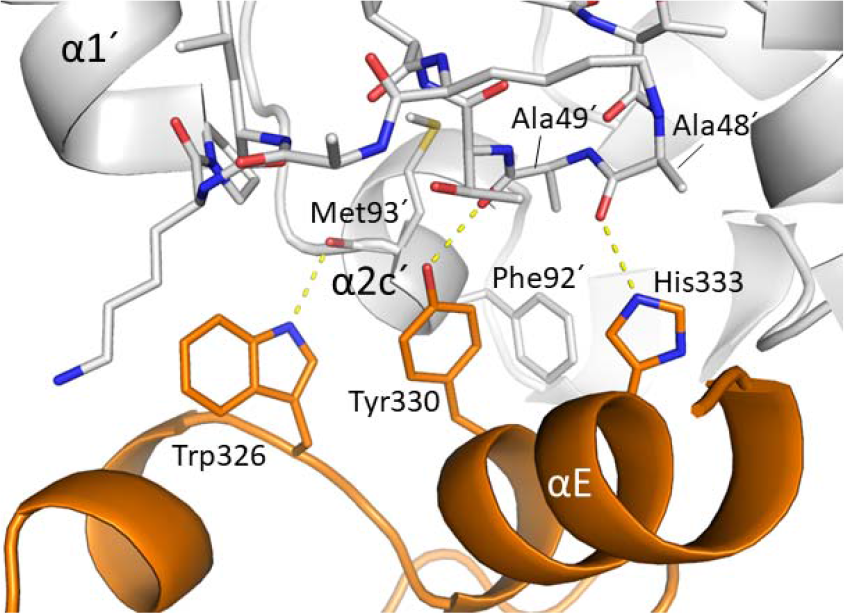
Aromatic cluster within the homodimer interface of *Z. mobilis* Tgt. The course of the main chain is mainly shown as ribbon. The two subunits are colored orange and grey, respectively. The side chains of Trp326, Tyr330, His333 and Phe92’ are shown as sticks. Met93’ as well as loop β1’α1’ are completely shown in stick representation. H-bonds are shown as dashed yellow lines.

The side chains of Trp326, Tyr330 and His333 are locked into position via their hydrogenbond donor functions, which form stable hydrogen bonds (H-bonds) to main chain carbonyl groups of residues within α-helix 2c’ and loop β1’α1’. Together with the succeeding α-helix 1, loop β1α1 forms an essential component of the dimer interface, referred to as “loop-helix motif”. In the crystal structures of Tgt featuring an intact dimer contact, this motif adopts a well-conserved conformation shielding the interface from water access. The binding of a dimer-destabilizing spiking ligand, however, mostly enforces a significantly different conformation of the loop-helix motif, which may culminate in its complete collapse.^18–20^ This phenomenon is also observed in several crystal structures of Tgt variants which contain mutations aimed at disturbing the architecture of the dimer interface.^21–23^

The primary goal of the present study was to investigate the importance of the H-bonds that Trp326, Tyr330 and His333 form to main chain carbonyl groups within the dimer mate. Therefore, we generated mutated variants in which one of these aromatic residues is mutated to phenylalanine, each, since the side chain of this residue is devoid of any H-bond donor function. We then analyzed the impact of the respective amino-acid exchange via X-ray crystallography, native mass spectrometry, thermal shift assay and enzyme kinetics. In addition, we analyzed a Tgt variant in which Trp95 is changed to phenylalanine, since we had gained evidence that this residue is involved in the stabilization of the loop-helix motif and, thus, of the homodimer interface as well.

## Results and discussion

### Structural and biochemical characterization of Tgt(Tyr330Phe)

Tyr330 of *Z. mobilis* Tgt, forming an H-bond to the main chain carbonyl oxygen of Ala49’ via its phenolic hydroxyl group, is strictly conserved in bacterial Tgt. To investigate the importance of this H-bond on the stability of the Tgt homodimer on a structural level we mutated Tyr330 to phenylalanine. In former work dealing with residues establishing the Tgt dimer interface, we had determined two crystal structures of a Lys52Met variant of Tgt, namely one of a crystal grown at pH 5.5 and one of a crystal grown at pH 8.5. Notably, the dimer destabilizing effect of the Lys52Met mutation, in particular in terms of structural changes within the loop-helix motif (Thr47 to Thr62), appears much more pronounced in a crystal grown at pH 5.5 than at pH 8.5.^21–22^ This prompted us to determine both a pH 5.5 and a pH 8.5 crystal structure of Tgt(Tyr330Phe) as well.

Indeed, disregarding the missing H-bond between the hydroxyl group of Tyr330 and Ala49’, virtually no changes in the architecture of the dimer interface are visible in the structure of Tgt(Tyr330Phe) determined from a crystal grown at pH 8.5. Thus, the loop-helix motif, which is well defined in the electron density map, adopts the same conformation as in wild type Tgt (Figure S1). In stark contrast, a considerable influence of the Tyr330Phe mutation on the loop-helix motif appears in the structure of this variant determined from a crystal grown at pH 5.5. Here, the first two residues of the loop-helix motif, Thr47 and Ala48, as well as the preceding Gly46 take a course which is completely different from that normally observed in crystal structures of Tgt. The remaining portion of the loop-helix motif, Ala49 to Thr62, obviously adopts no defined conformation or scatters over multiple conformations, as it is crystallographically not resolved (Figure S2). The collapse of the loop-helix motif is accompanied by a visible tilting of α-helices E and F entailing a shift of one subunit against the other one by circa 3 Å (Figure 2). These changes in subunit arrangement lead, compared with conventional *Z. mobilis* Tgt crystals, to measurably altered unit cell parameters. While the a-axis is shortened by approximately 6 Å, the β-angle is reduced by more than 2°. Notably, the same changes in unit cell parameters are observed for the above mentioned crystal of Tgt(Lys52Met) grown at pH 5.5^22^ as well as for several crystals of wild type Tgt resulting from co-crystallization of the enzyme with dimer interface-disturbing “spiking ligands”.^18–20^ It has to be emphasized that, in the absence of a spiking ligand, crystal growth at pH 5.5 has no consequence on unit cell parameters and on the conformation of the loop-helix motif in the case of wild type Tgt.

**Figure 2:**
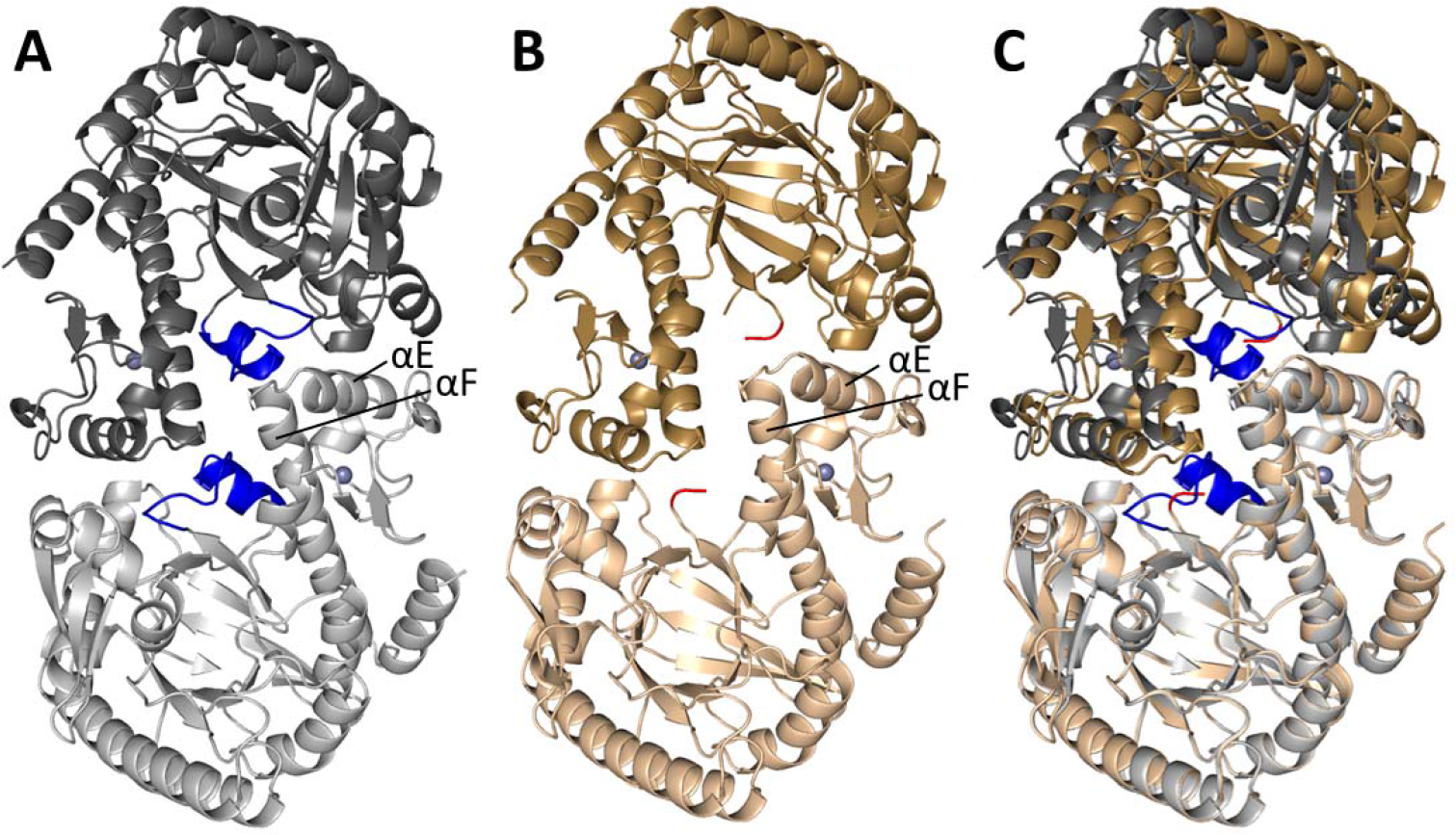
Ribbon representation of the *Z. mobilis* Tgt(Tyr330Phe) homodimer (A) crystallized at pH 8.5 and (B) crystallized at pH 5.5. (C) Superposition of (A) and (B). Subunits are indicated by different shades of color. The loop-helix motif (loop β1α1 and helix α1) is shown in blue and red, respectively. Zn^2+^-ions are colored grey.

Native mass spectrometry analysis performed under non-denaturing conditions confirmed the dimer destabilizing effect of the conservative amino acid exchange observed in the pH 5.5 crystal structure of Tgt(Tyr330Phe). At subunit concentrations of 10 μmol·L^−1^ and 1 μmol·L^−1^, the monomer portion of the mutated variant amounts to 45 ± 2% and 65 ± 5%, respectively (Figure 3). The determination of the Michaelis-Menten parameters of Tgt(Tyr330Phe) revealed that the introduced mutation does not influence the affinity of the enzyme to the tRNA substrate. Yet, it causes a slightly reduced catalytic activity since *k*_cat_ of Tgt(Tyr330Phe) is, compared with wild type Tgt, decreased by a factor of three (Table 1). To evaluate if the loss of the phenolic hydroxyl group in Tgt(Tyr330Phe) exerts influence on the overall stability of the enzyme we measured the “melting temperature” (*T*_m_) of this variant using a thermal shift assay.^24^ Under the adjusted conditions it amounts to 66.0 ± 1.5 °C which is slightly lower than that of wild type Tgt (*T*_m_ = 69.2 ± 0.3 °C) (Table 2).

**Figure 3:**
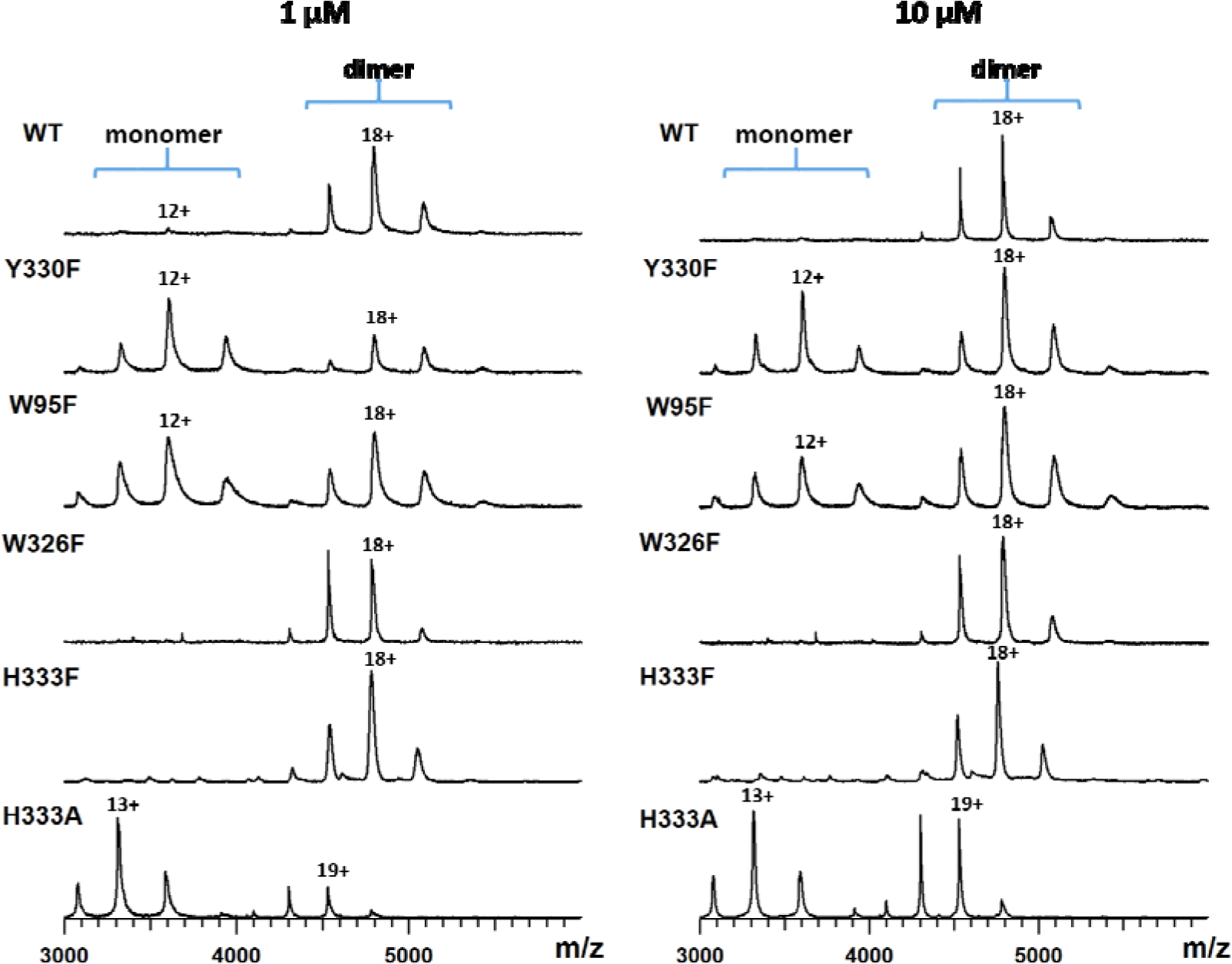
Native mass spectra of wild-type Tgt, Tgt(Tyr330Phe), Tgt(Trp95Phe), Tgt(Trp326Phe), Tgt(His333Phe), and Tgt(His333Ala) at 10 μmol·L^−1^ and 1 μmol·L^−1^ subunit concentration.

**Table 1.**
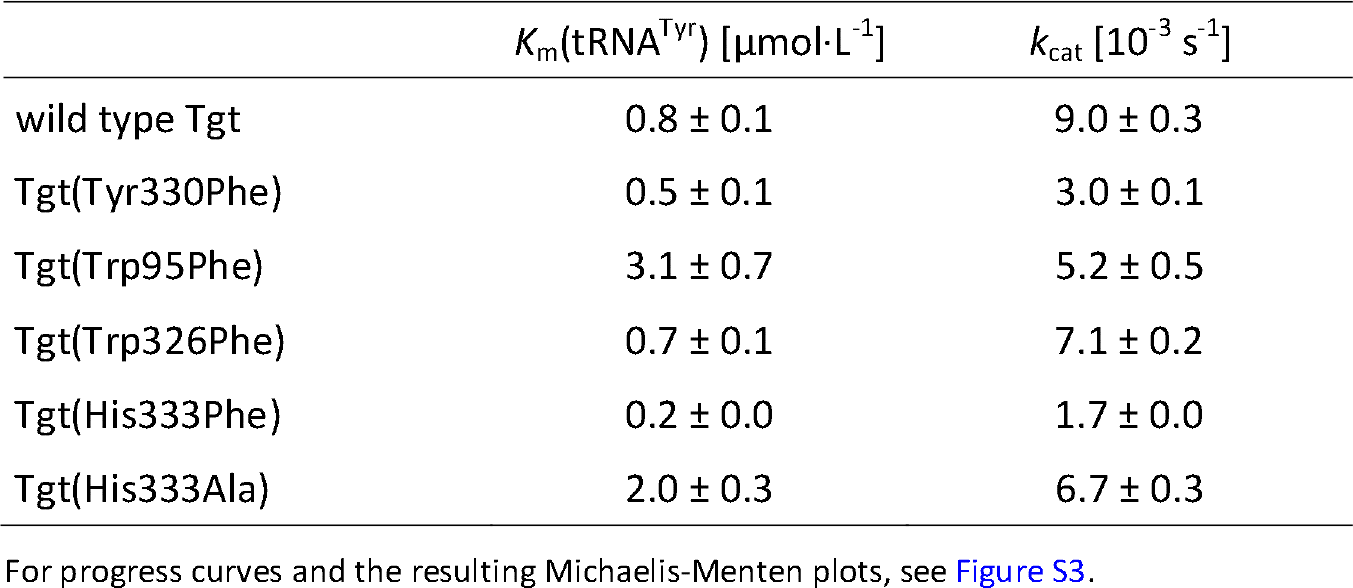
Kinetic parameters of Tgt variants determined in this study

**Table 2.** “Melting temperatures” (*T*_m_) of Tgt variants considered in this study

| | $T_m$ [°C] |
| --- | --- |
| wild type Tgt | $69.2 \pm 0.3$ |
| Tgt(Tyr330Phe) | $66.0 \pm 1.5$ |
| Tgt(Trp95Phe) | $53.7 \pm 0.5$ |
| Tgt(Trp326Phe)* | $67.8 \pm 0.4$ |
| Tgt(His333Phe) | $71.7 \pm 0.3$ |
| Tgt(His333Ala) | $63.7 \pm 0.0$ |

For melting curves and first derivatives thereof, see Figure S4.
\*The melting curve of Tgt(Trp326Phe) contains two inflection points indicating that unfolding occurs in two steps for this variant. Here, $T_m$ is defined as the midpoint temperature of protein unfolding transition. The temperatures at the minima of the first derivatives of the melting curve of Tgt(Trp326Phe) amount to $63.4 \pm 0.0$ °C and $73.7 \pm 0.1$ °C, respectively.

### The impact of Trp95Phe mutation on the Tgt homodimer interface

Also in crystal structures of [5-^19^F]-tryptophan-labeled Tgt(Trp95Phe) we observed a strikingly different impact of the introduced mutation on the dimer interface depending on the pH applied for crystallization. We had created this variant in the context of a project investigating the influence of certain Tgt ligands on the formation of the unproductive “twisted Tgt dimer”. To this end, we had labeled *Z. mobilis* Tgt with [5-^19^F]-tryptophan for NMR analyses. To facilitate the assignment of peaks within the obtained NMR spectra to particular [5-^19^F]-tryptophan residues we had generated mutated variants, in which individual tryptophan residues were changed to phenylalanine. With the intention to exclude major structural changes within these variants, we crystallized the obtained [5-^19^F]-tryptophan-labeled variants as well as [5-^19^F]-tryptophan-labeled wild type Tgt at pH 5.5 and determined their structures. As expected, the crystal structure of wild type Tgt labeled with [5-^19^F]-tryptophan did not reveal any significant structural changes compared with that of the unlabeled enzyme.

Unexpectedly, the unit cell of a crystal of [5-^19^F]-Trp-Tgt(Trp95Phe) showed a similar shortening of the a-axis by roughly 6 Å and reduction of the β-angle by some 2° as the crystal of Tgt(Tyr330Phe) grown at pH 5.5. Although Trp95 is in the wild type in close vicinity to the dimer interface, it is not directly involved in its formation. Rather, its side chain points toward the protein interior where it forms Van-der-Waals interactions with the side chains of Pro56, Val59 and Arg60,which are all components of α-helix 1 within the loop-helix motif. In addition, it forms hydrophobic contacts to Ile67, Met93, Arg97 and Ile99 as well as an H-bond to the main chain carbonyl oxygen of Pro98 (Figure 4). These interactions obviously contribute to the stable geometry of the loop-helix motif since the pH 5.5 crystal structure of [5-^19^F]-Trp-Tgt(Trp95Phe), which particularly lacks the ability to form an H-bond to the Pro98 carbonyl group, displays the same conformational changes as the pH 5.5 crystal structure of Tgt(Tyr330Phe). Gly46, Thr47 and Ala48 are dislocated while the portion of the loop-helix motif encompassing Ala49 to Thr62 has collapsed and is not visible in the electron density map. In contrast, no influence of the introduced mutation on the dimer interface is apparent in the pH 8.5 crystal structure of this variant. Clearly, compared with the indole moiety of Trp95, the smaller phenyl residue of Phe95 shows a reduced pattern of hydrophobic interactions and is not able to form an H-bond to the main chain carbonyl oxygen of Pro98. Nonetheless, the loop-helix motif as well as surrounding structural elements are in perfect congruence with those present in the crystal structure of wild type Tgt (Figure 4).

**Figure 4:**
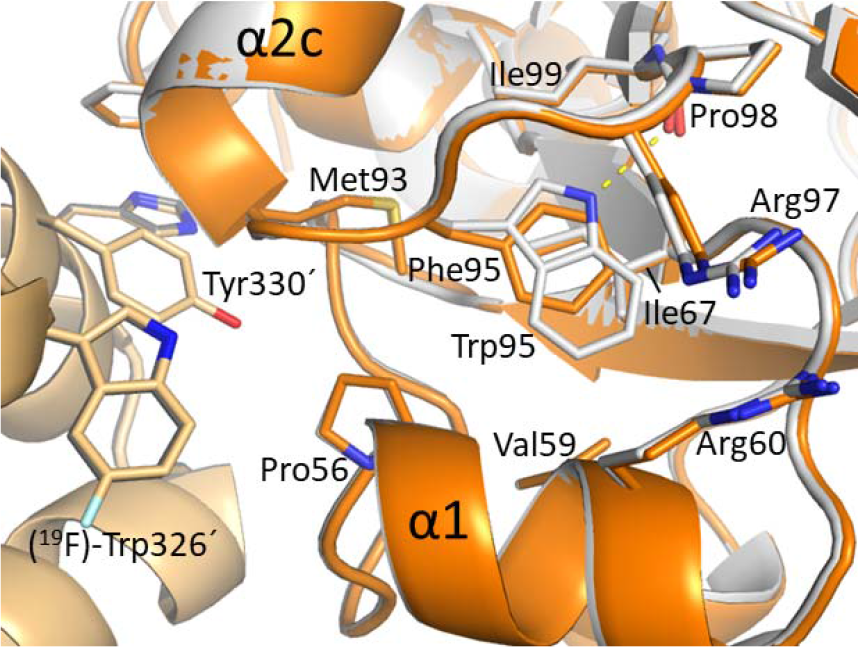
Superposition of wild type Tgt (carbon atoms grey and dark grey) and [5-^19^F]-Trp-Tgt(Trp95Phe) crystallized at pH 8.5 (carbon atoms orange and bright orange). The detail shows ribbon representations with relevant residues in stick representation. An H-bond formed between the indole moiety of Trp95 and the main chain carbonyl oxygen of Pro98 is indicated by a yellow dotted line.

The reduced dimer stability observed in the pH 5.5 crystal structure of Tgt(Trp95Phe) was verified by native mass spectrometry. At a subunit concentration of 10 μmol·L^−1^ the monomer portion of this variant amounts to 36 ± 3%, while it is increased to 51 ± 3% at a subunit concentrations of 1 μmol·L^−1^ (Figure 3). To find out if the reduced dimer stability caused by the Trp95Phe mutation is accompanied by a decrease in catalytic efficiency, we determined Michaelis-Menten parameters of this variant. While *k*_cat_ is slightly decreased, *K*_m_(tRNA^Tyr^) is marginally increased, with both changes being at the border of significance (Table 1). However, as shown via thermal shift assay, the substitution of Trp95 by phenylalanine leads to a strongly reduced thermal stability of the enzyme as the *T*_m_ of Tgt(Trp95Phe) is, compared with that of wild type Tgt, decreased by 15.5 °C (Table 2).

### The mutation of Trp326 to phenylalanine has no significant consequence on the architecture and stability of the homodimer interface

We also crystallized a [5-^19^F]-Trp-labeled Tgt variant in which Trp326 is mutated to phenylalanine. Via its indole moiety, Trp326 forms a stable H-bond to the main chain carbonyl oxygen of Met93’ in wild type *Z. mobilis* Tgt. Yet, in most of all bacterial Tgt enzymes Trp326 is replaced by phenylalanine or tyrosine. In line with this fact, the mutation of Trp326 to phenylalanine in *Z. mobilis* Tgt obviously has no detrimental effect on the geometry of the homodimer. No change in the architecture of its interface is observed in the pH 5.5 crystal structure of [5-^19^F]-Trp-Tgt(Trp326Phe) with the loop-helix motif being perfectly congruent with that observed in the crystal structures of ligand-free wild type Tgt (Figure S5). Furthermore, native mass spectrometry showed that the stability of the homodimer is not compromised upon mutation of Trp326 by phenylalanine. The mass spectra of this Tgt variant at 10 and 1 μmol·L^−1^ subunit concentration closely resemble those obtained from wild type Tgt with hardly any monomer being detectable even at the lower concentration (Figure 3).

Moreover, the mutation does not affect catalytic activity as reflected by *K*_cat_ and *K*_m_(tRNA) of Tgt(Trp326Phe), which do not significantly differ from that of wild type Tgt (Table 1). In addition, the introduced mutation does not seem to impair the thermal stability of the enzyme. However, the melting curve gained from Tgt(Trp326Phe) differs somewhat in shape from that normally obtained from Tgt as it contains two inflection points indicating that unfolding occurs in two steps (see Figure S4). Defining the melting temperature as the midpoint temperature of protein unfolding transition, *T*_m_ amounts to 67.8 ± 0.4 °C.

### The consequence of changing His333 to phenylalanine or alanine

Via its ε-NH group, the highly conserved His333 forms an H-bond to Ala48’ of the dimer mate. To determine the importance of the His333 imidazole for dimer stability, we mutated this residue to phenylalanine, whose aromatic side chain lacks any hydrogen-bond donor property.

The replacement of His333 by phenylalanine takes only moderate effect on dimer stability. Solely at a subunit concentration of 1 μmol·L^−1^, a low fraction of monomer (some 4%) is detectable for this variant, while at 10 μmol·L^−1^ subunit concentration no significant amount of monomer is present (Figure 3). In addition, the mutation has no harmful consequence on the thermal stability of the enzyme as the *T*_m_ of Tgt(His333Phe) is, compared with that of the wild type enzyme, even slightly increased (Table 2). Nonetheless, this variant shows a significantly reduced turnover number which is five-fold lower than that of wild type Tgt (Table 1).

In accordance with the low impact of the His333Phe mutation on dimer stability, the pH 8.5 crystal structure of Tgt(His333Phe) reveals no influence of the introduced mutation on the architecture of the dimer interface. Thus, the loop-helix motif is excellently defined in the electron density map and present in the same conformation as in the crystal structures of ligand-free wild type Tgt. Unexpectedly, no disturbance of the homodimer interface is visible in the pH 5.5 crystal structure of Tgt(His333Phe) either. Here, just like in the pH 8.5 crystal structure of this variant, the loop-helix motif is perfectly defined in the electron density map and its conformation does not differ from that in wild type Tgt (Figure S6). Consequently, there are no changes in unit cell parameters such as shortening of the a-axis and reduction of the β-angle which are observed in the pH 5.5 crystal structures of further Tgt variants harboring mutations that destabilize the dimer interface. Even though the His333Phe mutation has only moderate influence on dimer stability, this finding seemed surprising since, as mentioned above, the side chain imidazole of His333 forms an H-bond to Ala48’ within loop β1’α1’. Since this H-bond is clearly involved in stabilizing the conformation of loop β1’α1’ we had expected that its loss causes at least some structural alterations in this region. Ultimately, crystallization at pH 5.5 has resulted in the complete collapse of the loop-helix motif in further Tgt variants which still contain a histidine at position 333 and show a destabilized dimer interface. This finding raised the question if it might be the protonation of His333 at pH 5.5 that creates an electrostatic environment which severely affects the stability and conformation of the loop-helix motif, particularly of mutated variants that bear an additional destabilizing exchange (Tyr330Phe, Tyr95Phe, Lys52Met) or are affected by the binding of a dimer-destabilizing spiking ligand.

To corroborate this assumption, we changed His333 to alanine. Not only does the loss of the aromatic side chain result in the loss of favorable Van-der-Waals interactions with nearby hydrophobic residues but is also believed to facilitate the access of water into the dimer interface. Accordingly, we expected this mutation to severely affect homodimer stability. Yet, just as the side chain of phenylalanine the side chain methyl group of the introduced alanine is unable to accept a proton.

Native mass spectrometry showed that, actually, the absence of the aromatic residue in Tgt(His333Ala) entails a pronounced destabilization of the homodimer. At a subunit concentration of 10 μmol·L^−1^, the monomer portion of this variant amounts to 45%, while at a subunit concentration of 1 μmol·L^−1^ it increases to 80% (Figure 3). In spite of the strongly decreased stability of the Tgt(His333Ala) homodimer, only moderate impact of the introduced mutation is observed in the pH 8.5 crystal structure of this variant. The lack of the imidazole does not cause any change in the conformation of the remaining residues of the aromatic cluster. In addition, the course of the loop-helix motif is nearly identical to that observed in crystal structures of wild type Tgt. Merely Thr47 to Ala49 do obviously not adopt a fixed conformation, since they are ill-defined in the electron density map, which is why these residues were omitted from the final model deposited with the protein data bank (pdb-code: 6h7c). Notably, the same applies to the structure of Tgt(His333Ala) determined from a crystal that was grown at pH 5.5. Also here, the loop-helix motif is found in its usual conformation, apart from residues Thr47 to Ala49, which are ill-defined in the electron density map (Figure S7). Obviously, histidine at position 333 plays a very special role regarding the integrity of the dimer interface. According to its p*K*_a_ in aqueous solution, His333 presumably adopts a proton at pH 5.5. As various crystal structures show, the positively charged histidine of this residue triggers the collapse of the loop-helix motif in Tgt variants whose dimer stability is, due to a mutation, compared with the wild-type protein, reduced. Supposedly, the latter is sufficiently stable to tolerate a positive charge in the area of the aromatic cluster without structural consequences. The mechanism underlying this phenomenon and the putative role of His333 for the functional properties of the enzyme are, however, unclear at present since our crystal structures do not provide an obvious explanation. Nevertheless, this shows the importance of a well-balanced composition of hydrophobic aromatic residues in the cluster, which can easily be perturbed by the introduction of a charge. Remarkably, once this charge is removed (even by the introduction of a residue like alanine, which has clearly a detrimental effect on the stability of the aromatic cluster) the overall-integrity of the loop-helix motif is rescued.

The mutation of His333 to alanine causes a markedly reduced thermal stability of the enzyme since the *T*_m_ of Tgt(His333Ala) is, compared with that of the wild type enzyme, decreased by 5.5 °C. This does, however, not involve a significant reduction of catalytic activity since the turnover number of this variant is hardly lower than that of wild type Tgt (Table 1). Importantly, we were able to reproduce the unexpectedly high *k*_cat_ of Tgt(His333Ala) with a different preparation of this mutated variant without, however, repeating the cost-intensive complete determination of Michaelis-Menten parameters (data not shown).

Notably, in early studies from 1995 and 1996, Chong et al.^25^ and Garcia et al.^26^ mutated His317 of *Escherichia coli* Tgt, which corresponds to His333 of *Z. mobilis* Tgt, to alanine and cysteine, respectively. Via native polyacrylamide gel electrophoresis (PAGE) this group had shown in a preceding work, that the enzyme forms a homo-oligomer.^27^ Its stoichiometry, however, was still unknown, since no crystal structure of a Tgt enzyme was available at the time these studies were conducted. After visual inspection of the primary structure of *E. coli* Tgt, Chong et al.^25^ rightly assumed that the enzyme contains a Zn^2+^ ion, which is coordinated by the sulfhydryl groups of three cysteines and the imidazole residue of one histidine. While they correctly identified Cys302, Cys304 and Cys307 (corresponding to Cys318, Cys320 und Cys323 in *Z. mobilis* Tgt) as the Zn^2+^-coordinating cysteines, they erroneously assumed His317 to be the Zn^2+^-coordinating histidine. To investigate the putative role of His317 in Zn^2+^ binding they mutated this residue to alanine. While *E. coli* Tgt(His317Ala) still contains Zn^2+^, the authors showed via native PAGE that its ability to form a homo-oligomer and its capacity to bind substrate tRNA are significantly impaired. Consequently (and inconsistent with our results), they observed reduced catalytic activity for *E. coli* Tgt(His317Ala) without, however, determining Michaelis-Menten parameters for this mutated variant. In a subsequent study, Garcia et al.^26^ mutated His317 of *E. coli* Tgt to cysteine. Via native PAGE they showed that, under the conditions tested, also Tgt(His317Cys) does not form a homo-oligomer in the absence of substrate tRNA. In the presence of substrate tRNA, however, it clearly forms the intact active complex, which is meanwhile known to consist of the Tgt homodimer and one tRNA molecule. Indeed, similar to *Z. mobilis* Tgt(His333Ala), *k*_cat_ and *K*_m_(tRNA) of *E. coli* Tgt(His317Cys) are, compared to the wild type enzyme, hardly compromised.

### The Tgt homodimer is highly persistent as shown by native mass spectrometry

The plasmid we use for the recombinant expression of the *Z. mobilis tgt* gene encodes the Tgt enzyme fused to an N-terminal *Strep*-tag II^®^, which may be proteolytically removed by means of a thrombin cleavage site.^21,23^ To gain an estimate of the constancy of the Tgt homodimer we used this construct to analyze the kinetics of subunit exchange. For this purpose, we mixed Tgt containing the N-terminal Strep-tag II^®^ with an equimolar amount of Tgt lacking this tag. By means of noncovalent mass spectrometry, we then monitored the appearance of “heterodimeric” Tgt consisting of both a tagged and an untagged subunit in addition to Tgt consisting of only one subunit species, respectively. While, immediately (2 min) after combining the two preparations, virtually no “heterodimeric” Tgt was identified in mass spectra, equilibrium was reached after more than 10 h. In contrast, equilibrium was reached in less than 10 min in the case of Tgt(His333Ala), the variant generated in this study exhibiting the most strongly impaired dimer stability (Figure 5).

**Figure 5:**
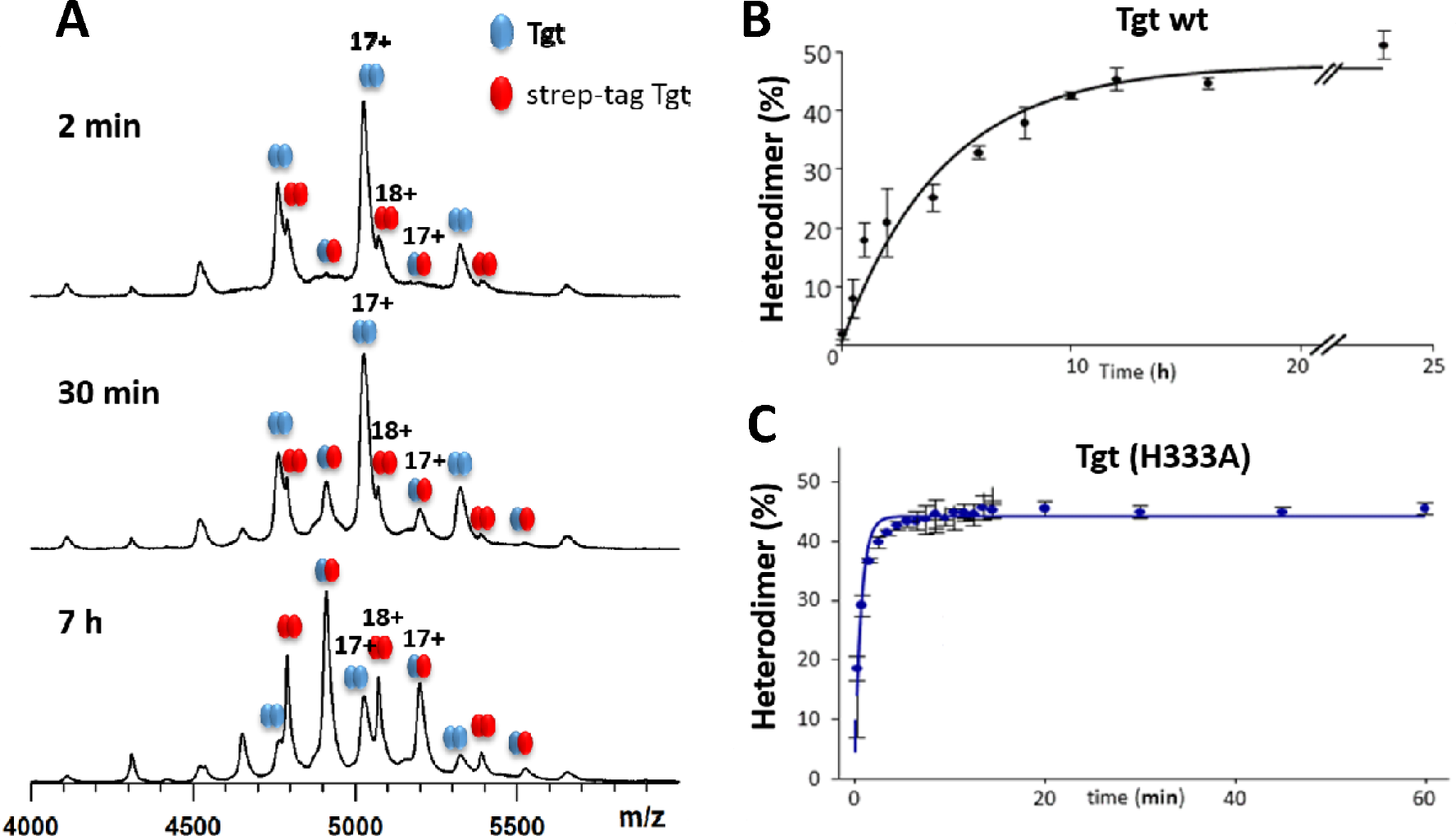
(A) Spectra of time resolved native MS during analysis of subunit exchange (wild type Tgt). (B) Kinetics of subunit exchange in wild type Tgt and (C) in Tgt(His333Ala).

## Conclusions

In the present study we have investigated the importance of the H-bonds that Trp326, Tyr330, and His333 form to residues of the dimer mate for the homodimer stability of *Z. mobilis* Tgt (Figure 1). For this purpose, we created Tgt variants in which one of the named residues is mutated to phenlylalanine, each. In conformance with the fact that in most bacterial Tgt enzymes Trp326 is replaced by phenylalanine or tyrosine, the substitution of the tryptophan indole by a phenyl residue has no significant consequences on homodimer stability, overall stability and catalytic efficiency of the enzyme. In contrast, the loss of the H-bond caused by the mutation of the strictly conserved Tyr330 to phenylalanine leads to strongly reduced homodimer formation concomitant with slightly decreased thermal stability and three-fold lowered enzymatic activity. Ultimately, His333 is nearly invariant in bacterial Tgt enzymes of known sequence (solely in the Tgt of *Blochmannia floridanus* it is replaced by tyrosine). Yet, the replacement of this residue by phenylalanine results in only slight reduction of homodimer stability with no decrease in overall stability being observable. However, the His333Phe mutation adversely influences catalytic activity since *k*_cat_ of Tgt(His333Phe) is five-fold reduced compared to wild type Tgt.

Both Tyr330 and His333 form H-bonds to main chain carbonyl groups of residues within loop β1’α1’ of the loop-helix motif, which efficiently shields the dimer interface from water access. In the course of a different study, we had accidentally gained evidence that Trp95, which makes numerous interactions with residues of the loop-helix motif, including an H-bond to the Pro98 carbonyl group, is required for the stabilization of this structural element. Accordingly, we included the structural and biochemical characterization of Tgt(Trp95Phe) into our study. We showed that the substitution of Trp95 by phenylalanine not only results in strongly reduced homodimer formation but also in a massive loss of overall stability since *T*_m_ of Tgt(Trp95Phe) is decreased by some 15 °C compared with wild type Tgt. In contrast, the mutation hardly entails any loss of catalytic activity.

Remarkably, the dimer destabilizing effects of the Trp95Phe as well as of the Tyr330Phe mutations only become visible in crystal structures that have been determined from crystals grown at pH 5.5, but not from crystals grown at pH 8.5. While in the pH 5.5 crystal structures of both variants the loop-helix motif has collapsed and is almost completely invisible in the electron density map, virtually no changes in the architecture of the dimer interface are observed in the corresponding pH 8.5 crystal structures. The same phenomenon had been observed before for Tgt(Lys52Met), a variant created and analyzed in preceding studies.^21,22^ Obviously, slightly acidic pH has a destabilizing effect on the loop-helix motif and leads to its collapse in the presence of a mutation that further destabilizes this structural element. In the pH 5.5 crystal structure of Tgt(His333Phe), however, no consequence of the introduced mutation is observed, although it causes the loss of an H-bond to the loop-helix motif of the dimer mate. This brought us to speculate that the protonation of the His333 imidazole, which cannot take place in Tgt(His333Phe), might have a detrimental effect on the stability of the loop-helix motif. To confirm this assumption we changed His333 to alanine, which sacrifices a large portion of hydrophobic contacts within the aromatic cluster, but cannot accept a proton as well. Indeed, although the introduced mutation results in severe destabilization of the homodimer, no collapse of the loop-helix motif is observed in the pH 5.5 crystal structure of Tgt(His333Ala). Rather, it looks identical to the pH 8.5 crystal structure of this variant, where this motif is largely present in its usual conformation with only three residues being ill-defined in the electron density map. We take this as strong evidence that the protonation of the His333 imidazole intensifies the effect of mutations affecting dimer stability, which leads to the breakdown of the loop-helix motif in the pH 5.5 crystal structures of such Tgt variants.

In the present study as well as in preceding studies,^21,23^ we made the surprising observation that Tgt variants harboring mutations that severely influence the stability of the homodimer show only slight or almost no reduction in catalytic activity. Likely, the presence of a tRNA substrate strongly promotes the formation, maintenance and thus stability of the homodimer and, consequently, the functioning of the enzyme. Furthermore, the decrease of catalytic activity caused by a dimer interface-disturbing mutation does obviously not correlate with the severity of dimer destabilization. E.g., the mutation of His333 to alanine leads to considerable loss of dimer stability but to hardly any reduction in catalytic efficiency. In contrast, the mutation of His333 to phenylalanine only slightly impacts dimer stability but results in a turnover number which is five-fold reduced compared to wild type Tgt. Obviously, mutations of residues involved in dimer formation also tend to affect the nearby catalytic center and/or substrate binding site, but to different extent. Thus, in a former study, we had determined a turnover number which is decreased by more than one order of magnitude for the already mentioned Tgt(Lys52Met) variant, although the introduced mutation has only moderate effect on dimer stability.^22^ The crystal structure of Tgt in complex with an RNA substrate shows that Lys52 not only forms a salt bridge to Glu339’ of the dimer mate, but also makes an extensive Van-der-Waals contact to uracil35 of the bound tRNA molecule.^16^ Presumably, this contact is important for correct substrate orientation and, as a consequence, for efficient catalysis. Similar, although less evident reasons may account for the reduced turnover numbers measured for Tgt(His333Phe) and further Tgt variants with decreased dimer stability.

## Methods

### Mutagenesis, protein production and purification

Overexpression of the wild type *tgt* gene and mutated variants thereof as well as protein purification were done as described by Jakobi et al.^21^ The production of [5-^19^F]-tryptophan-labeled Tgt will be described elsewhere. For DNA manipulation, plasmids were prepared using the peqGOLD plasmid miniprep kit (PEQLAB, Erlangen, Germany). Site directed mutagenesis was performed via the QuikChange Lightning kit (Agilent, Santa Clara, CA, USA) according to the vendor’s instructions. The required DNA primers were purchased from Eurofins MWG Operon (Ebersberg, Germany). In each case, the complete tgt gene was re-sequenced (by Eurofins MWG Operon) to confirm both the presence of the desired mutation and the absence of any further, inadvertent mutation.

### Preparation of tRNA^Tyr^

Unmodified *E. coli* tRNA^Tyr^ (ECY2)^28^ required for kinetic analyses was synthesized via in vitro transcription by T7-RNA-Polymerase. The reaction mixture (20 mmol·L^−1^ MgCl_2_, 80 mmol·L^−1^ HEPES pH 7.5, 1 mmol·L^−1^ spermidine, 5 mmol·L^−1^ DTT, 0.05 U pyrophosphatase, 30 μg linearised DNA template, 3.75 mmol·L^−1^ of each NTP, 5% DMSO and 1 μmol·L^−1^ T7-RNA polymerase) was incubated for 4 h at 37 °C. The transcription product was subjected to phenol/chloroform extraction using an equal volume of (1:1) acid-phenol:chloroform (pH 4.5) and precipitated by the addition of 0.1 volume of 3 mol·L^−1^ sodium acetate (pH 5.2) and 2 volumes of ethanol. After centrifugation (10 min at 16,500 × *g* at 4 °C) the supernatant was discarded. The pellet was dried, dissolved in ddH_2_O and subjected to preparative denaturing (8 mol·L^−1^ urea) 8% polyacrylamide gel electrophoresis. The RNA band of interest was detected by UV shadowing and excised from the gel using a sterile scalpel. Subsequently, the tRNA was eluted from the gel slice overnight under shaking at 900 rpm using a thermomixer (Eppendorf, Hamburg, Germany) in 1 mol·L^−1^ sodium acetate (pH 5.2) at 4 °C. Finally, the purified tRNA was ethanol precipitated as described above and, after drying, re-dissolved in ddH_2_O. The concentration of tRNA was determined via UV photometry (λ = 280 nm) using a NanoDrop™ 2000c spectrophotometer (Thermo Fisher Scientific, Waltham, MA, USA).

### Kinetic characterisation of Tgt variants

The determination of *k_ca_* and *K*_m_(tRNA^Tyr^) of *Z. mobilis* Tgt variants was done by monitoring the insertion of [8-^3^H]-guanine (1.5 Ci·mmol^−1^, American Radiolabeled Chemicals, Inc., Saint Louis, MO, USA) into tRNA as described by Biela et al..^29^ In each case, the enzyme was used at a subunit concentration of 150 nmol·L^−1^. While the concentration of [^3^H]-labeled guanine was kept constant at 10 μmol·L^−1^, the concentration of tRNA^Tyr^ varied between 0.26 μmol·L^−1^ and 15 μmol·L^−1^. Kinetic parameters were calculated using the programme *GraphPad PRISM* (version 7.04).

### Thermal shift assay

Protein unfolding within a temperature range between 10 °C and 100 °C in 320 increments was monitored via the fluorescence of SYPRO™ orange (Thermo Fisher Scientific, Waltham, MA, USA) essentially as described by Niesen et al..^24^ Thermal shift experiments were done in triplicate using a QuantStudio 3 Real-Time PCR System (Thermo Fisher Scientific) equipped with a MicroAmp^®^ Fast 96-Well Reaction Plate (0.1 mL) (Thermo Fischer Scientific). The volume of each sample was 20 μL with 24 μg of protein being diluted in buffer containing 10 mmol·L^−1^ TrisHCl, pH 7.8, 1 mmol·L^−1^ EDTA, 1 mmol·L^−1^ DTT, 2 mol·L^−1^ NaCl and 2 × SYPRO orange dye. The wavelengths for excitation and emission were 470 ± 15 nm and 575 ± 15 nm, respectively. Data were processed with Excel. The minimum of the first derivative of each melting curve was used to determine melting temperature, *T*_m_.

### Native mass spectrometry experiments

A hybrid Q-IM-TOF mass spectrometer (Synapt G2, Waters, Manchester, UK) coupled to an automated chip-based nanoelectospray device (Triversa Nanomate, Advion, Ithaca, USA) operating in the positive mode was used to perform native mass spectrometry experiments. The pressure of the nebulizer gas and capillary voltage were set to 0.6 psi and 1.75 kV respectively. The backing pressure of the instrument was fixed to 6 mbar and the cone voltage was set to 80 V to ensure ion transmission and reduce dimer dissociation. Calibration of the mass spectrometer was performed from monovalent ions produced by a 2 mgom?^1^ cesium iodide solution in 1:1 2-propanol/water (v/v). Prior to native mass spectrometry analysis, all the enzymes were buffer exchanged against 1 mol·L^−1^ ammonium acetate at pH 7.5 using microcentrifuge gel-filtration columns (Zeba 0.5 mL, Thermo Scientific, Rockford, IL). Protein concentration was determined with a NanoDrop™ spectrophotometer (Thermo Fisher Scientific, France). Proteins were diluted to obtain a final concentration of 1 and 10 μmol·L^−1^ to study the monomer/dimer ratio, or to 2.5 μmol·L^−1^ for time-resolved native mass spectromery experiments. In the latter experiments, equimolar mixtures of wild type Tgt and Tgt(His333Ala) with their corresponding streptavidin-labeled variants at 2.5 μmol·L^−1^ concentration were used to study subunit exchange kinetics. The heterodimer/homodimer ratio of streptavidin-labeled and unlabeled proteins were calculated from their relative mass intensities at different incubation times (from 0 min to 25 h). Mass spectrometry data interpretation was performed using MassLynx 4.1 (Waters).

### Crystallization, data collection, structure determination, model building and refinement

All Tgt variants were crystallized using the hanging-drop vapor diffusion method at 18 °C. 1 μL of protein solution (ca. 12 g·L^−1^ in 10 mmol·L^−1^ TrisHCl pH 7.8, 1 mmol·L^−1^ EDTA, 2 mol·L^−1^ NaCl) was mixed with 1 μL of reservoir solution (100 mmol·L^−1^ TrisHCl pH 8.5 or 100 mmol·L^−1^ MES pH 5.5, 10% (v/v) DMSO, 13% (w/v) PEG 8000). Crystals grew after a few days in the presence of 650 μL reservoir solution. Prior to data collection, crystals were transferred to a buffer containing 50 mmol·L^−1^ TrisHCl pH 8.5 or 50 mmol·L^−1^ MES pH 5.5, 300 mmol·L^−1^ NaCl, 2% (v/v) DMSO, 4% (w/v) PEG 8000 and 30% (v/v) glycerol as cryo-protectant for a few seconds and vitrified in liquid nitrogen.

Diffraction data were collected at 100 K at the synchrotron beamlines and wavelengths that are listed in Table S1. Diffraction data from Tgt(His333Ala) (pH 5.5) and Tgt(His333Phe) (pH 5.5) were collected in house on an Incoatec IμS operated at 45 kV and 650 μA using copper wavelength (1.54178Å) and equipped with a Mar-dtb and Mar345 image plate detector. All diffraction images were indexed, processed and scaled using XDS^30^ and *XDSAPP^31^* Data collection and refinement statistics are summarized in Table S1. All structures were determined via molecular replacement using the program *Phaser^32^* from the *CCP4* suite.^33^ PDB entry 1pud served as search model. Model building was done in *Coot*,^34^ while the program *Phenix^35^* was used for structure refinement.

### Figure preparation

Structural figures were prepared using *Pymol* (http://www.pymol.org).

### Protein data bank accession codes

Coordinates and structure factors have been deposited under following accession codes:

Tgt(Tyr330Phe) pH 5.5: 6yfx; Tgt(Tyr330Phe) pH 8.5: 6ygk; Tgt(His333Ala) pH 5.5: 6yry;

Tgt(His333Ala) pH 8.5: 6h7c; Tgt(His333Phe) pH 5.5: 6z0d; Tgt(His333Phe) pH 8.5: 6yfw; [5-^19^F]-Trp-Tgt(Trp95Phe) pH 5.5: 6ygm; [5-^19^F]-Trp-Tgt(Trp95Phe) pH 8.5: 6ygo; [5-^19^F]-Trp-Tgt(Trp326Phe) pH 5.5: 6ygl; [5-^19^F]-Trp-Tgt pH 5.5: 6ygp.

## Supporting information

Figures S1 – S7; Table S1

## Acknowledgements

We gratefully acknowledge the beamline staff at BESSY II (Helmholtz-Zentrum Berlin, Germany) for outstanding support during data collection and Christian Sohn for his help during in-house X-ray data collection.

## References

1. Tuorto, F., Legrand, C., Cirzi, C., Federico, G., Liebers, R., Müller, M., Ehrenhofer-Murray, A.E., Dittmar, G., Gröne, H.-J., and Lyko, F. (2018) Queuosine-modified tRNAs confer nutritional control of protein translation. EMBO J. 37, e99777.

2. Müller, M., Legrand, C., Tuorto, F., Kelly, V.P., Atlasi, Y., Lyko, F., and Ehrenhofer-Murray (2019) Queuine links translational control in eukaryotes to a micronutrient from bacteria. Nucleic Acids Res. 47, 3711–3727.

3. Phillips, G., Yacoubi, B.E., Lyons, B., Alvarez, S., Iwata-Reuyl, D., and de Crécy-Lagard, V. (2008) Biosynthesis of 7-deazaguanosine-modified tRNA nucleosides: a new role for GTP cyclohydrolase I. J. Bacteriol. 190, 7876–7884.

4. McCarty, R.M., Somogyi, Á., and Bandarian, V. (2009) *Escherichia coli* QueD is a 6-carboxy-5,6,7,8-tetrahydropterin synthase. Biochemistry 48, 2301–2303.

5. McCarty, R.M., Somogyi, Á., Lin, G., Jacobsen, N.E., and Bandarian, V. (2009) The deazapurine biosynthetic pathway revealed: in vitro enzymatic synthesis of preQ_0_ from guanosin *5’*-triphosphate in four steps. Biochemistry 48, 3847–3852.

6. McCarty, R.M., Krebs, C., and Bandarian, V. (2013) Spectroscopic, steady-state kinetic, and mechanistic characterization of the radical SAM enzyme QueE, which catalyzes a complex cyclization reaction in the biosynthesis of 7-deazapurines. Biochemistry 52, 188–198.

7. Van Lanen, S.G., Reader, J.S., Swairjo, M.A., de Crécy-Lagard, V., Lee, B., and Iwata-Reuyl, D. (2005) From cyclohydrolase to oxidoreductase: discovery of nitrile reductase activity in a common fold. Proc. Natl. Acad. Sci. USA 102, 4264–4269.

8. Okada, N. and Nishimura, S. (1979) Isolation and characterization of a guanine insertion enzyme, a specific tRNA transglycosylase, from *Escherichia coli*. J. Biol. Chem. 254, 3061–3066.

9. Grimm, C., Ficner, R., Sgraja, T., Haebel, P., Klebe, G., and Reuter, K. (2006) Crystal structure of *Bacillus subtilis* S-adenosylmethionine:tRNA ribosyl transferase-isomerase. Biochem. Biophys. Res. Commun. 351, 695–701.

10. Van Lanen, S.G., Kinzie, S.D., Matthew, S., Link, T., Culp, J. and Iwata-Reuyl, D. (2003) tRNA modification by S-adenosylmethionine:tRNA ribosyltransferase-isomerase. J. Biol. Chem. 278, 10491–10499.

11. Van Lanen, S.G. and Iwata-Reuyl, D. (2003) Kinetic mechanism of the tRNA modifying enzyme S-adenosylmethionine:tRNA ribosyltransferase-isomerase (QueA). Biochemistry 42, 5312–5320.

12. Miles, Z.D., McCarty, R.M., Molnar, G., and Bandarian, V. (2011) Discovery of epoxyqueuosine (oQ) reductase reveals parallels between halorespiration and tRNA modification. Proc. Natl. Acad. Sci. USA 108, 73268–7372.

13. Zallot, R., Ross, R., Chen, W. H., Bruner, S.D., Limbach, P.A., and de Crécy-Lagard, V. (2017) Identification of a novel epoxyqueuosine reductase family by comparative genomics. ACS Chem. Biol. 12, 844–851.

14. Durand, J.M.B., Okada, N., Tobe, T., Watarai, M., Fukuda, I., Suzuki, T., Nakata, N., Komatsu, K., Yoshikawa, M., and Sasakawa, C. (1994) *vacC*, a virulence-associated chromosomal locus of *Shigella flexneri*, is homologous to *tgt*, a gene encoding tRNA-guanine transglycosylase (Tgt) of *Escherichia coli* K-12. J. Bacteriol. 176, 4627–4634.

15. Romier, C., Reuter, K., Suck, D., and Ficner, R. (1996) Crystal structure of tRNA-guanine transglycosylase: RNA modification by base exchange. EMBO J. 15, 2850–2857.

16. Xie, W., Liu, X., and Huang, R.H. (2003) Chemical trapping and crystal structure analysis of a catalytic tRNA guanine transglycosylase covalent intermediate. Nature Struct. Biol. 10, 781–788.

17. Stengl, B., Meyer, E.A., Heine, A., Brenk, R., Diederich, F., and Klebe, G. (2007) Crystal structures of tRNA-guanine transglycosylase (TGT) in complex with novel and potent inhibitors unravel pronounced induced-fit adaptations and suggest dimer formation upon substrate binding. J. Mol. Biol. 370, 492–511.

18. Immekus, F., Barandun, L.J., Betz, M., Debaene, F., Petiot, S., Sanglier-Cianferani, S., Reuter, K., Diederich, F., and Klebe, G. (2013) Launching spiking ligands into a protein-protein interface: a promising strategy to destabilize and break interface formation in a tRNA modifying enzyme. ACS Chem. Biol. 8, 1163–1178.

19. Ehrmann, F.R., Stojko, J., Metz, A., Debaene, F., Barandun, L.J., Heine, A., Diederich, F., Cianférani, S., Reuter, K., and Klebe, G. (2017) Soaking suggests “alternative facts”: only cocrystallization discloses major ligand-induced interface rearrangements of a homodimeric tRNA-binding protein indicating a novel mode-of-inhibition. PLOS ONE 12, e0175723.

20. Ehrmann, F.R., Kalim, J., Pfaffeneder, T., Bernet, B., Hohn, C., Schäfer, E., Botzanowski, T., Cianférani, S., Heine, A., Reuter, K., Diederich, F., and Klebe, G. (2018) Swapping interface contacts in the homodimeric tRNA-guanine transglycosylase: an option for functional regulation. Angew. Chem. Int. Ed. 57, 10085–10090.

21. Jakobi, S., Nguyen, T.X.P., Debaene, F., Metz, A., Sanglier-Cianférani, S., Reuter, K., and Klebe, G. (2014) Hot-spot analysis to dissect the functional protein-protein interface of a tRNA-modifying enzyme. Proteins Struct. Funct. Bioinf. 82, 2713–2732.

22. Ritschel, T., Atmanene, C., Reuter, K., Van Dorsselaer, A., Sanglier-Cianferani. S, and Klebe, G. (2009) An integrative approach combining noncovalent mass spectrometry, enzyme kinetics and X-ray crystallography to decipher Tgt protein-protein and protein-RNA interaction. J. Mol. Biol. 393, 833–847.

23. Jakobi, S., Nguyen, T.X.P., Debaene, F., Cianférani, S., Reuter, K., and Klebe, G. (2015) What glues a homodimer together: systematic analysis of the stabilizing effect of an aromatic hot spot in the protein-protein interface of the tRNA-modifying enzyme Tgt. ACS Chem. Biol. 10, 1897–1907.

24. Niesen, F.H., Berglund, H., and Vevadi, M. (2007) The use of differential scanning fluorimetry to detect ligand interactions that promote protein stability. Nature Protocols 2, 2212–2221.

25. Chong, S., Curnow, A.W., and Garcia, G.A. (1995) tRNA-guanine transglycosylase from *Escherichia coli* is a zinc metalloprotein. Site-directed mutagenesis studies to identify the zinc ligands. Biochemistry 34, 3694–3701.

26. Garcia, G.A., Tierney, D.L., Chong, S., Clark, K., and Penner-Hahn, J.E. (1996) X-ray absorption spectroscopy of the Zinc site in tRNA-guanine transglycosylase from *Escherichia coli*. Biochemistry 35, 3133–3139.

27. Garcia, G.A., Koch, K.A., and Chong, S. (1993) tRNA-guanine transglycosylase from *Escherichia coli* - overexpression, purification and quaternary structure. J. Mol. Biol. 231, 489–497.

28. Curnow, A.W., Kung, F.-L., Koch, K.A., and Garcia, G.A. (1993) tRNA-guanine transglycosylase from *Escherichia coli:* gross tRNA structural requirements for recognition. Biochemistry 32, 5239–5246.

29. Biela, I., Tidten-Luksch, N., Immekus, F., Glinca, S., Nguyen, T.X.P., Gerber, H.-D., Heine, A., Klebe, G., and Reuter, K. (2013) Investigation of specificity determinants in bacterial tRNA-guanine transglycosylase reveals queuine, the substrate of its eucaryotic counterpart, as inhibitor. PLOS ONE 8, e64240.

30. Kabsch, W. (2010) XDS. Acta Crystallogr. D66, 125–132.

31. Krug, M., Weiss, M.S., Heinemann, U., and Mueller, U. (2012) XDSAPP: a graphical user interface for the convenient processing of diffraction data using XDS. J. Appl. Crystallogr. 45, 568–572.

32. McCoy, A.J., Grosse-Kunstleve, R.W., Adams, P.D., Winn, M.D., Storoni, L.C., and Read, R.J. (2007) Phaser crystallographic software. 40, 658–674.

33. Pottertin, E., Briggs, P., Turkenberg, M., and Dodson, E. (2003) A graphical interface to the CCP4 program suite. Acta Crystallogr. D59, 1131–1137.

34. Emsley, P., Lohkamp, B., Scott, W. G., and Cowtan, K. (2010) Features and development of Coot. Acta Crystallogr. D66, 486–501.

35. Adams, P.D., Afonine, P.V., Bunkóczi, G., Chen, V.B., Davis, I.W., Echols, N., Headd, J.J., Hung, L.-W., Kapral, G.J., Grosse-Kunstleve, R.W., McCoy, A.J., Moriarty, N.W., Oeffner, R., Read, R.J., Richardson, D.C., Richardson, J.S., Terwilliger, T.C., and Zwart, P.H. (2010) *PHENIX:* a comprehensive Python-based system for macromolecular structure solution. Acta Crystallogr. D66, 213–221.

36. Laskowski, R.A., MacArthur, M.W., Moss, D.S., and Thornton, J.M. (1993) PROCHECK: a program to check the stereochemical quality of protein structures. J. Appl. Cryst. 26, 283–291.

37. Kleywegt, G.J., Zou, J.Y., Kjeldgaard, M., and Jones, T.A. (2001) Around O, in: International Tables for Crystallography, Vol. F. Crystallography of Biological Macromolecules (Rossmann, M. G. & Arnold, E., Editors). Chapter 17.1, pp. 353-356, 366-367. Dordrecht: Kluwer Academic Publishers, The Netherlands.

