## Supplementary Figures and Tables for "The importance of charge in perturbing the aromatic glue stabilizing the protein-protein interface of homodimeric tRNA-guanine transglycosylase"

#### Figure S1

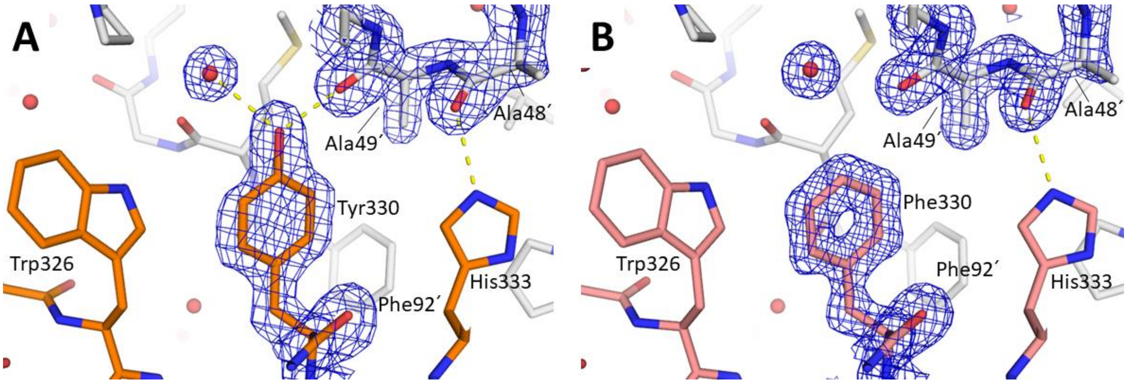

Detail of the homodimer interfaces of (A) wild type Tgt (pdb-code: 1p0d) and (B) Tgt(Tyr330Phe) crystallized at pH 8.5 (pdb-code: 6ygk) comprising the aromatic cluster as well as Ala48´ and Ala49´. Densities ($2f_o$-$f_c$) are contoured at 1.5 σ. H-bonds are shown as dashed yellow lines.

#### Figure S2

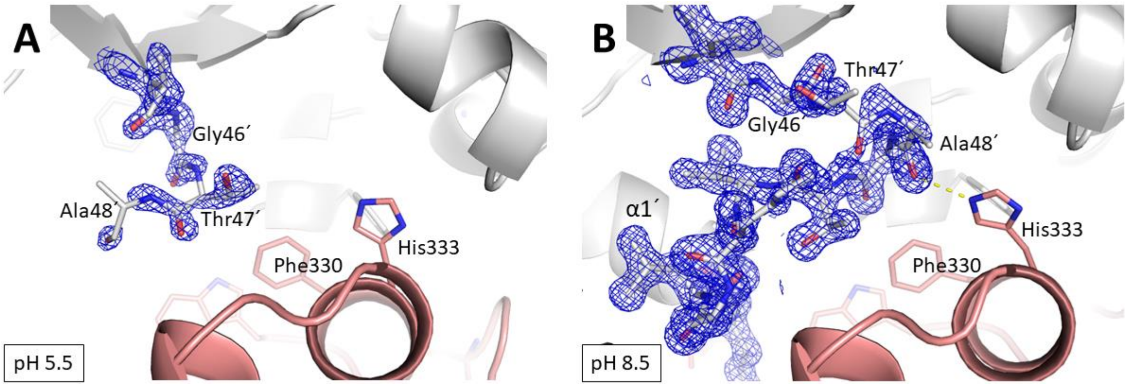

The loop-helix motif of (A) Tgt(Tyr330Phe) crystallized at pH 5.5 (pdb-code: 6yfx) and (B) Tgt(Tyr330Phe) crystallized at pH 8.5 (pdb-code: 6ygk). Densities ($2f_o$-$f_c$) are shown for residues of loop β1´α1´ and contoured at 1.5 σ. H-bonds are shown as dashed yellow lines.

### Figure S3

#### Wild type Tgt

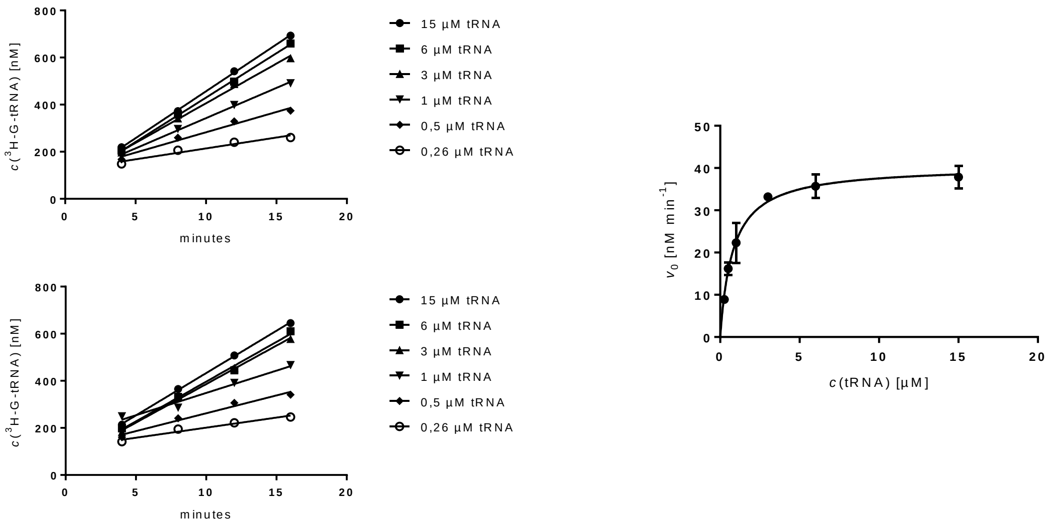

#### Tgt(Trp95Phe)

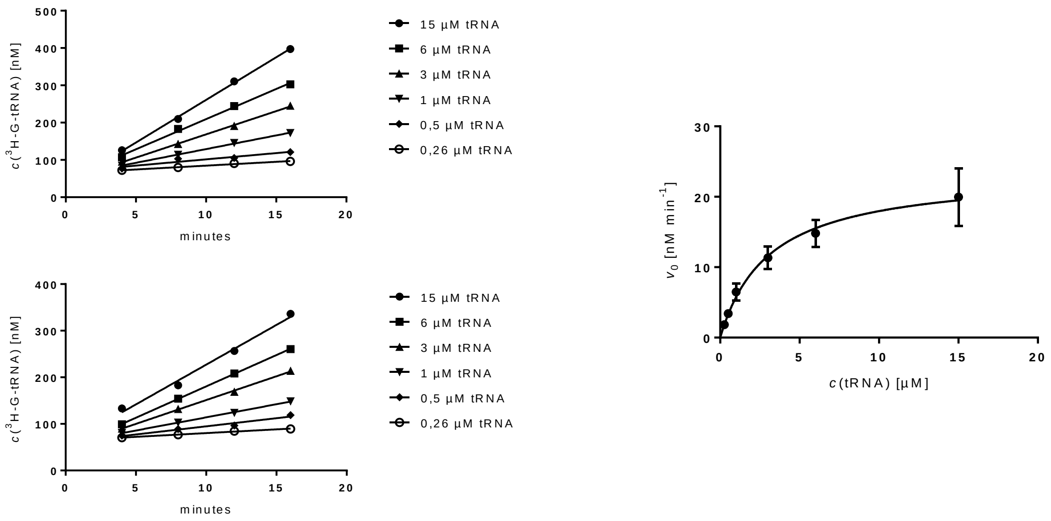

#### Tgt(Trp326Phe)

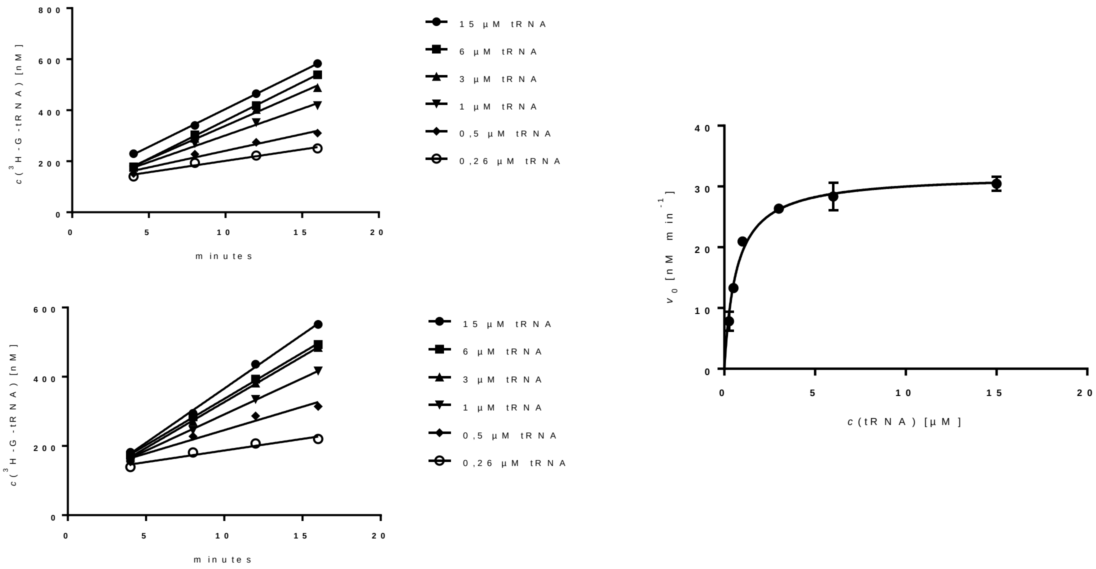

#### Tgt(Tyr330Phe)

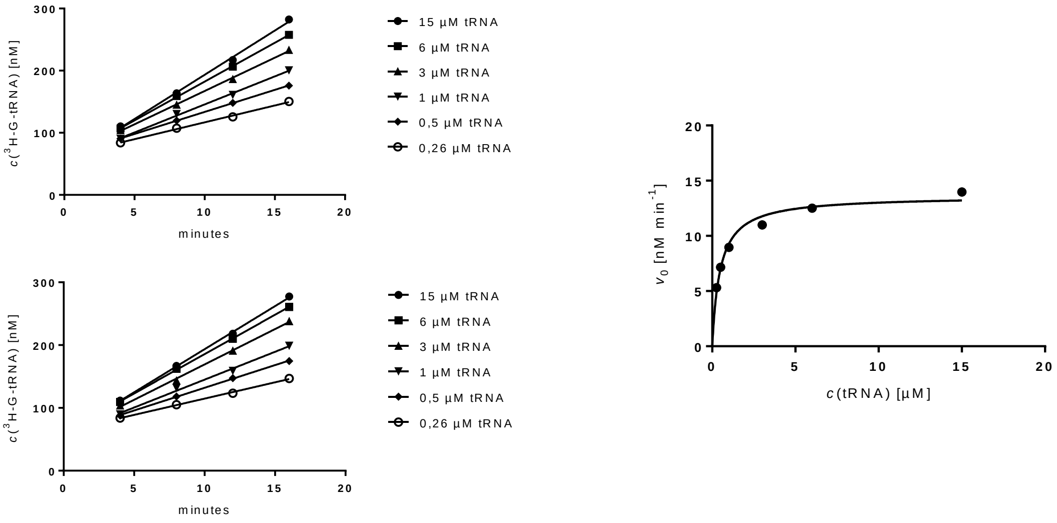

#### Tgt(His333Phe)

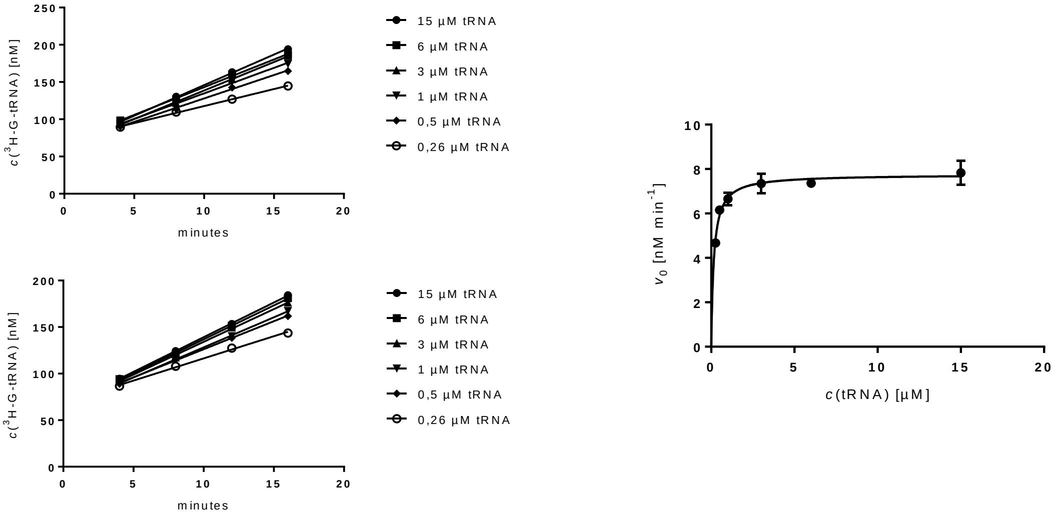

#### Tgt(His333Ala)

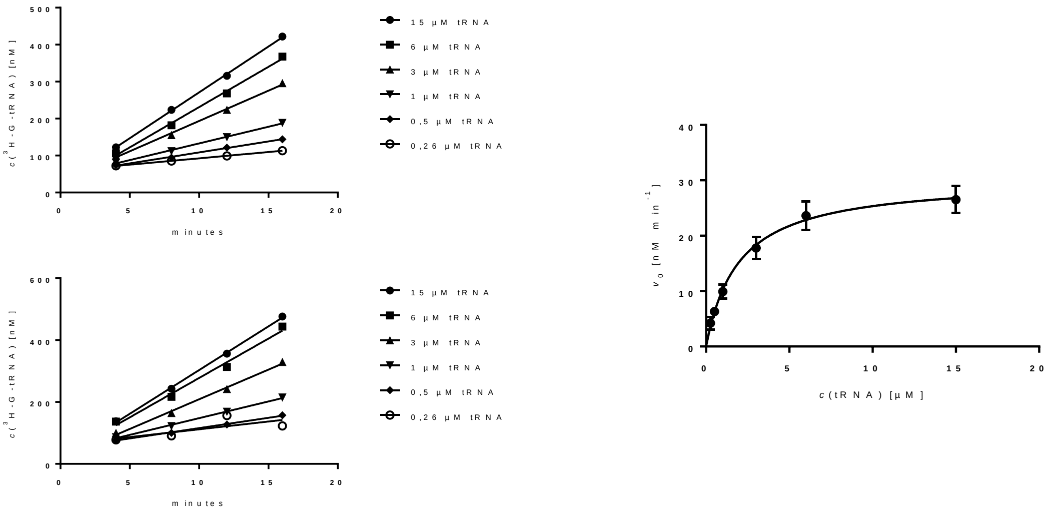

Progress curves and resulting Michaelis-Menten plots of *Zymomonas mobilis* Tgt and mutated Tgt variants thereof. Plots were generated by the program *GraphPad PRISM* (version 7.04).

### Figure S4

Wild type Tgt

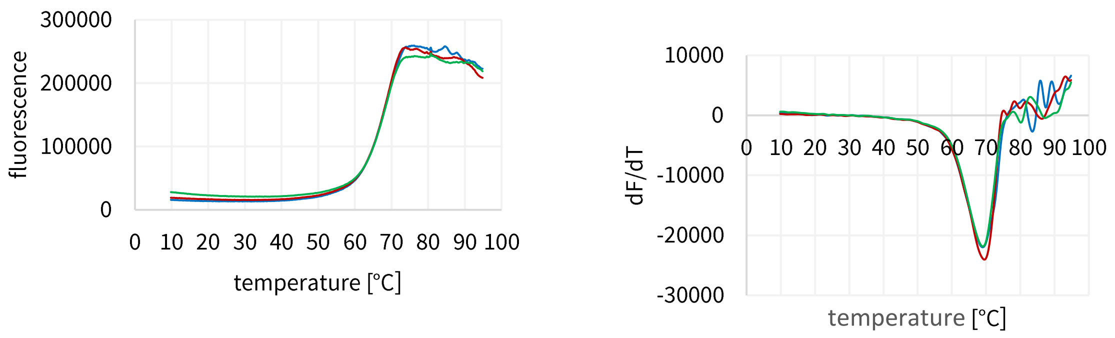

Tgt(Tyr330Phe)

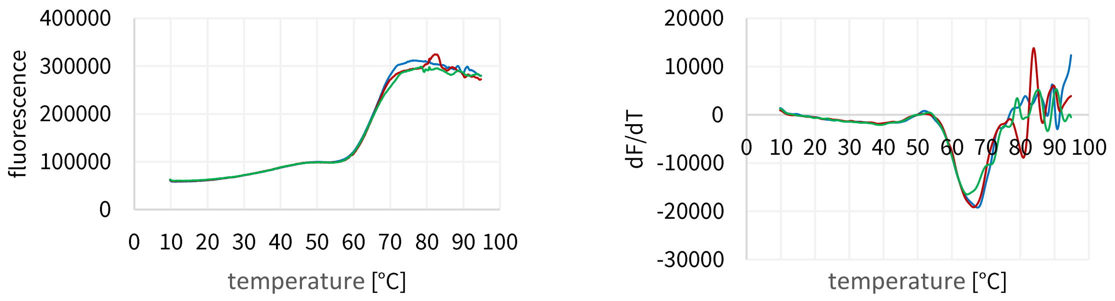

Tgt(Trp95Phe)

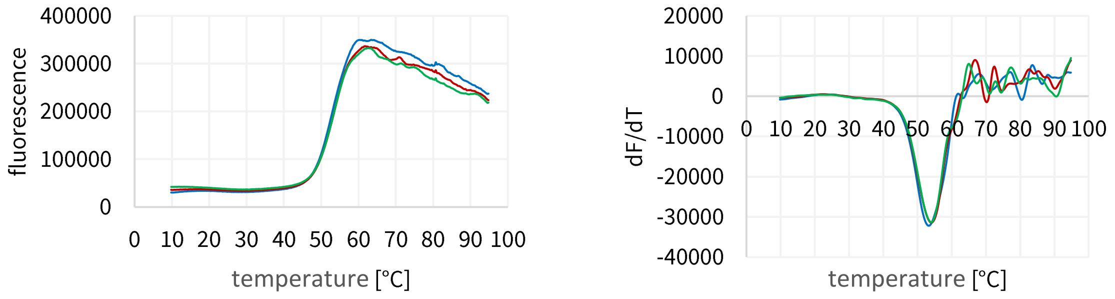

Tgt(Trp326Phe)

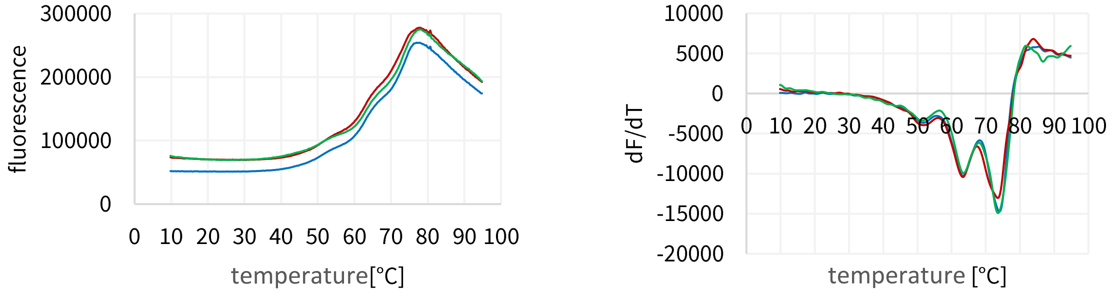

Tgt(His333Phe)

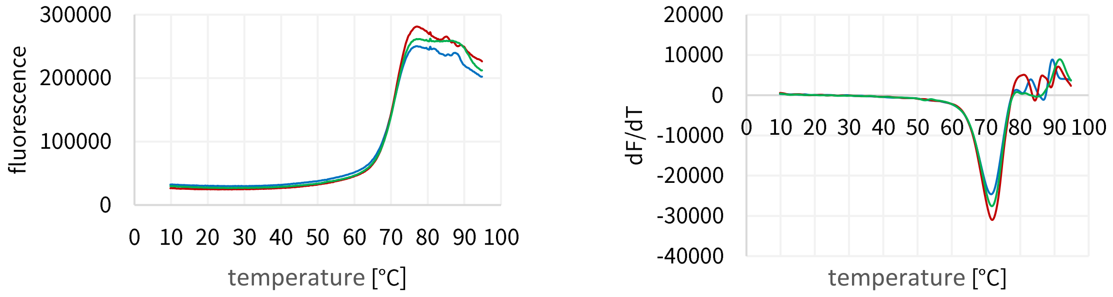

Tgt(His333Ala)

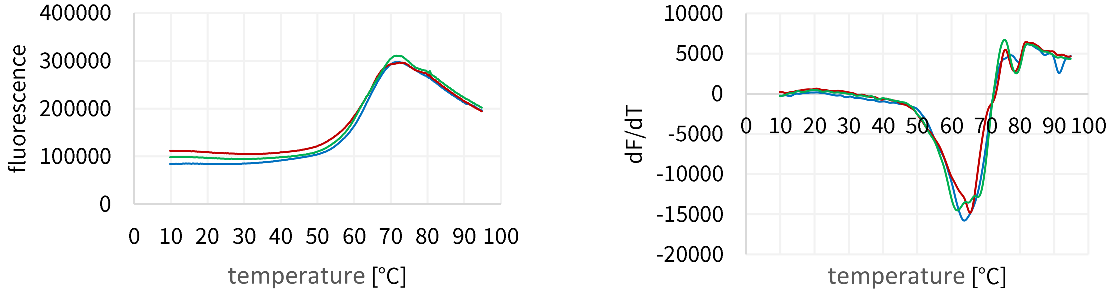

Melting curves and their first derivatives resulting from the thermal shift assay of *Z. mobilis* Tgt variants. Plots were created using the program *Excel*.

### Figure S5

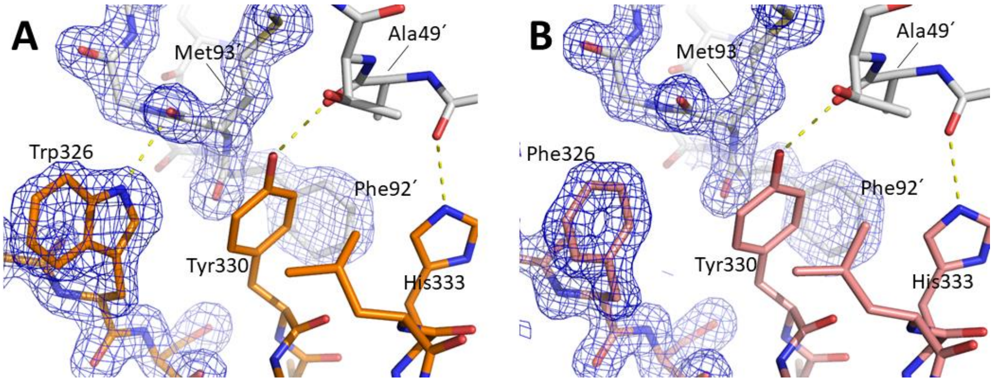

Detail of the homodimer interfaces of (A) wild type Tgt (pdb-code: 1p0d) and (B) Tgt(Trp326Phe) crystallized at pH 5.5 (pdb-code: 6ygl) comprising the aromatic cluster. Densities ($2f_o$-$f_c$) are contoured at 1.5 σ. H-bonds are shown as a dashed yellow lines.

### Figure S6

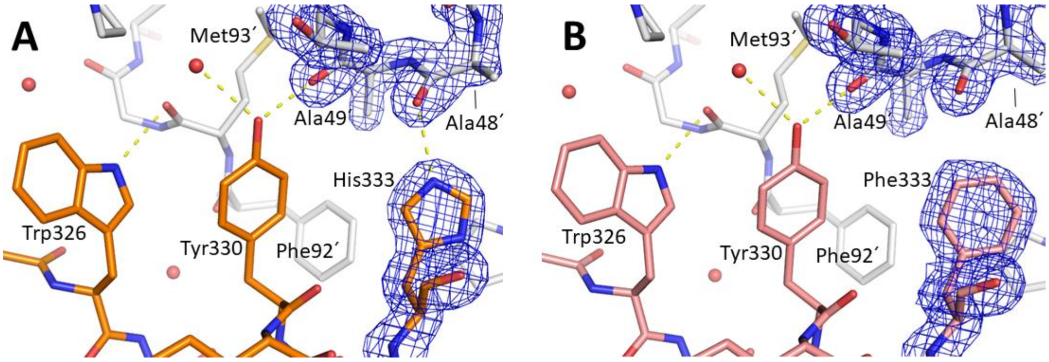

Detail of the homodimer interfaces of (A) wild type Tgt (pdb-code: 1p0d) and (B) Tgt(His333Phe) crystallized at pH 5.5 (pdb-code: 6z0d) comprising the aromatic cluster. Densities ($2f_o$-$f_c$) are contoured at 1.5 σ. H-bonds are shown as a dashed yellow lines.

### Figure S7

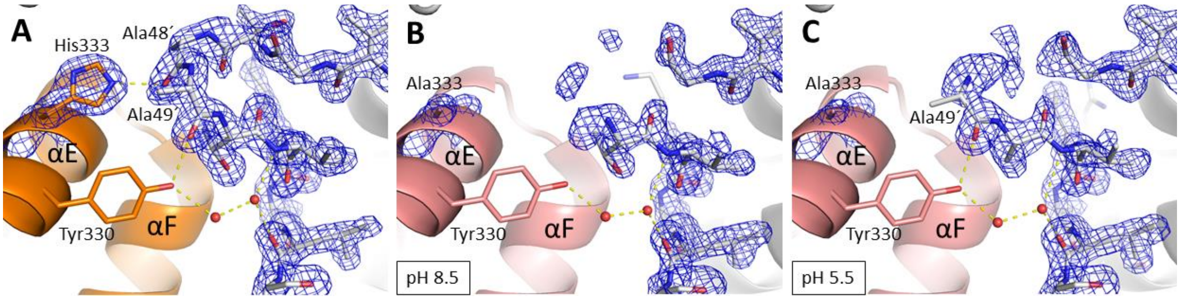

Detail of the homodimer interface of (A) wild type Tgt (pdb-code: 1p0d), (B) Tgt(His333Ala) crystallized at pH 8.5 (pdb-code: 6h7c) and (C) Tgt(His333Ala) crystallized at pH 5.5 (pdb-code: 6yry). Densities ($2f_o$-$f_c$) are contoured at 1.5 σ and shown for His/Ala333 as well as for Val45 to Lys55 within the loop-helix motif. H-bonds are shown as a dashed yellow lines.

### Table S1

#### Crystallographic data collection, processing and refinement statistics

##### Table S1a

| Crystal data | Tgt(Tyr330Phe) pH 5.5 | Tgt(Tyr330Phe) pH 8.5 | Tgt(His333Ala) pH 5.5 | Tgt(His333Ala) pH 8.5 |
|---|---|---|---|---|
| **PDB ID** | **6yfx** | **6ygk** | **6yry** | **6h7c** |
| ***(A) Data collection and processing*** | | | | |
| Collection site | BESSY 14.1 | BESSY 14.1 | "in house" | BESSY 14.2 |
| Wavelength [Å] | 0.91841 | 0.91841 | 1.54178 | 0.91841 |
| *Unit cell parameters* | | | | |
| Space group | *C*2 | *C*2 | *C*2 | *C*2 |
| *a, b, c* [Å] | 84.6, 65.1, 71.4 | 90.8, 65.0, 70.3 | 90.3, 65.2, 70.9 | 91.3, 65.0, 70.5 |
| β [°] | 93.7 | 96.0 | 96.5 | 95.9 |
| ***(B) Diffraction data*** | | | | |
| Resolution range [Å] | 42.56-1.38 (1.46-1.38) | 43.40-1.40 (1.48-1.40) | 44.87-1.82 (1.93-1.82) | 35.00-1.68 (1.72-1.68) |
| No. of unique reflections[a] | 79276 (12689) | 77939 (11334) | 35506 (5627) | 46188 (2556) |
| $R(I)_{sym}$[a] [%] | 5.0 (49.7) | 3.5 (35.7) | 7.1 (48.7) | 6.8 (31.7) |
| Completeness [%] | 98.9 (98.5) | 97.1 (87.9) | 97.1 (95.7) | 98.8 (94.4) |
| Multiplicity | 4.3 (4.3) | 3.3 (3.0) | 2.9 (2.8) | 3.0 (2.8) |
| Mean $I/\sigma(I)$ | 14.9 (2.5) | 18.0 (2.7) | 9.4 (2.7) | 13.0 (3.5) |
| Matthews coefficient [$Å^3$/Da] | 2.3 | 2.4 | 2.4 | 2.5 |
| Solvent content [%] | 46.0 | 48.8 | 49.0 | 50.0 |
| Overall *B* factor from Wilson Plot [$Å^2$] | 15.7 | 15.0 | 23.4 | 16.4 |
| No. proteins per asymmetric unit | 1 | 1 | 1 | 1 |
| ***(C) Refinement with Phenix* (version 1.16_3549)** | | | | |
| Resolution range [Å] | 42.56 - 1.38 | 43.40 – 1.40 | 44.87 – 1.82 | 26.47 - 1.68 |
| Reflections used in refinement | 75312 / 3964 | 74042 / 3897 | 33730 / 1776 | 43830 / 2331 |
| $R_{work}$[b]/$R_{free}$[c] [%] | 14.1 / 16.3 | 12.9 / 15.4 | 17.5 / 20.4 | 16.0 / 18.4 |
| *No. of atoms (non-hydrogen)* | | | | |
| Protein | 2784 | 2968 | 2788 | 2823 |
| water molecules | 279 | 357 | 274 | 270 |
| $Zn^{2+}$ | 1 | 1 | 1 | 1 |
| Glycerol | 6 | 24 | 18 | 30 |
| *rmsd from ideality* | | | | |
| bond angles [°] | 0.9 | 0.9 | 0.8 | 0.9 |
| bond length [Å] | 0.007 | 0.007 | 0.006 | 0.007 |
| *Ramachandran plot*[d] | | | | |
| most favored [%] | 94.7 | 94.7 | 95.4 | 96.5 |
| additionally allowed [%] | 5.0 | 5.0 | 4.2 | 3.2 |
| generously allowed [%] | 0.3 | 0.3 | 0.3 | 0.3 |
| *Mean B factor*[e] *[$Å^2$]* | | | | |
| Protein atoms | 22.3 | 20.4 | 25.8 | 20.9 |
| Water molecules | 32.0 | 31.2 | 33.4 | 32.8 |
| $Zn^{2+}$ | 18.9 | 13.6 | 22.7 | 14.8 |
| Glycerol atoms | 25.9 | 32.8 | 33.9 | 35.8 |

#### Table S1b

| Crystal data | Tgt(His333Phe) pH 5.5 | Tgt(His333Phe) pH 8.5 | $^{19}$F-Trp-Tgt(Trp95Phe) pH 5.5 | $^{19}$F-Trp-Tgt(Trp95Phe) pH 8.5 |
|---|---|---|---|---|
| **PDB ID** | **6z0d** | **6yfw** | **6ygm** | **6ygo** |
| ***(A) Data collection and processing*** | | | | |
| Collection site | "in house" | BESSY 14.1 | BESSY 14.1 | BESSY 14.1 |
| Wavelength [Å] | 1.54178 | 0.91841 | 0.91841 | 0.91841 |
| *Unit cell parameters* | | | | |
| Space group | *C*2 | *C*2 | *C*2 | *C*2 |
| *a, b, c* [Å] | 90.8, 65.3, 70.6 | 90.6, 65.0, 70.5 | 84.5, 65.0, 71.3 | 90.8, 64.9, 70.8 |
| β [°] | 96.3 | 95.9 | 94.0 | 96.0 |
| ***(B) Diffraction data*** | | | | |
| Resolution range [Å] | 40.00-1.65 (1.75-1.65) | 43.43-1.26 (1.34-1.26) | 42.56-1.23 (1.31-1.23) | 43.49-1.26 (1.34-1.26) |
| No. of unique reflections[a] | 47815 (6664) | 106813 (17052) | 107419 (16748) | 109116 (17332) |
| $R(I)_{sym}$[a] [%] | 3.6 (13.2) | 4.6 (45.3) | 3.8 (49.9) | 4.5 (49.3) |
| Completeness [%] | 96.7 (83.9) | 98.0 (97.2) | 97.5 (94.6) | 98.7 (97.2) |
| Multiplicity | 3.6 (3.3) | 3.0 (3.0) | 3.8 (3.7) | 3.8 (3.6) |
| Mean $I/\sigma(I)$ | 22.3 (7.7) | 11.9 (2.0) | 16.1 (2.4) | 14.2 (2.1) |
| Matthews coefficient [$Å^3$/Da] | 2.6 | 2.4 | 2.3 | 2.4 |
| Solvent content [%] | 52.5 | 48.8 | 45.7 | 48.9 |
| Overall *B* factor from Wilson Plot [$Å^2$] | 15.2 | 14.1 | 14.3 | 14.1 |
| No. proteins per asymmetric unit | 1 | 1 | 1 | 1 |
| ***(C) Refinement with Phenix* (version 1.16_3549)** | | | | |
| Resolution range [Å] | 43.6 – 1.65 | 43.40 – 1.26 | 42.17 - 1.23 | 43.49 - 1.26 |
| Reflections used in refinement | 45424 / 2391 | 101466 / 5341 | 102047 / 5371 | 103658 / 5456 |
| $R_{work}$[b]/$R_{free}$[c] [%] | 16.4 / 19.4 | 13.0 / 15.4 | 13.6 / 16.0 | 12.2 / 14.5 |
| *No. of atoms (non-hydrogen)* | | | | |
| Protein | 2867 | 2984 | 2786 | 2965 |
| water molecules | 380 | 361 | 334 | 354 |
| $Zn^{2+}$ | 1 | 1 | 1 | 1 |
| Glycerol | 6 | 24 | - | 24 |
| *rmsd from ideality* | | | | |
| bond angles [°] | 0.8 | 0.9 | 0.9 | 1.0 |
| bond length [Å] | 0.006 | 0.007 | 0.007 | 0.007 |
| *Ramachandran plot*[d] | | | | |
| most favored [%] | 94.4 | 95.9 | 95.7 | 95.0 |
| additionally allowed [%] | 5.2 | 3.8 | 4.0 | 4.7 |
| generously allowed [%] | 0.3 | 0.3 | 0.3 | 0.3 |
| *Mean B factor*[e] *[$Å^2$]* | | | | |
| Protein atoms | 19.0 | 19.5 | 21.4 | 20.1 |
| Water molecules | 29.4 | 33.2 | 35 | 34.4 |
| $Zn^{2+}$ | 16.6 | 12.5 | 16.2 | 12.8 |
| Glycerol atoms | 26.7 | 29.9 | - | 33.2 |

#### Table S1c

| Crystal data | $^{19}$F-Trp-Tgt(Trp326Phe) pH 5.5 | $^{19}$F-Trp-Tgt pH 5.5 |
|---|---|---|
| **PDB ID** | **6ygl** | **6ygp** |
| ***(A) Data collection and processing*** | | |
| Collection site | BESSY 14.1 | BESSY 14.1 |
| Wavelength [Å] | 0.91841 | 0.91841 |
| *Unit cell parameters* | | |
| Space group | *C*2 | *C*2 |
| *a, b, c* [Å] | 91.1, 65.2, 70.7 | 90.8, 65.2, 70.6 |
| β [°] | 96.1 | 96.1 |
| ***(B) Diffraction data*** | | |
| Resolution range [Å] | 45.31-1.48 (1.57-1.48) | 45.16-1.33 (1.41-1.33) |
| No. of unique reflections[a] | 66504 (10258) | 90420 (13633) |
| $R(I)_{sym}^{a}$ [%] | 3.9 (43.6) | 3.2 (36.7) |
| Completeness [%] | 97.4 (93.4) | 96.2 (90.3) |
| Multiplicity | 2.8 (2.7) | 3.1 (2.9) |
| Mean $I/\sigma(I)$ | 15.1 (2.0) | 17.0 (2.4) |
| Matthews coefficient [$Å^3$/Da] | 2.4 | 2.4 |
| Solvent content [%] | 49.3 | 48.9 |
| Overall *B* factor from Wilson Plot [$Å^2$] | 18.1 | 15.7 |
| No. proteins per asymmetric unit | 1 | 1 |
| ***(C) Refinement with Phenix*** **(version 1.16_3549)** | | |
| Resolution range [Å] | 45.31 - 1.48 | 45.2 – 1.33 |
| Reflections used in refinement | 63177 / 3326 | 85899 / 4521 |
| $R_{work}^{b}/R_{free}^{c}$ [%] | 13.2 / 16.4 | 12.1 / 14.8 |
| *No. of atoms (non-hydrogen)* | | |
| Protein | 2860 | 2909 |
| water molecules | 351 | 377 |
| $Zn^{2+}$ | 1 | 1 |
| Glycerol | 12 | 18 |
| *rmsd from ideality* | | |
| bond angles [°] | 1.0 | 1.0 |
| bond length [Å] | 0.007 | 0.007 |
| *Ramachandran plot*[d] | | |
| most favored [%] | 95.8 | 95.8 |
| additionally allowed [%] | 3.8 | 3.8 |
| generously allowed [%] | 0.3 | 0.3 |
| *Mean B factor*[e] *[$Å^2$]* | | |
| Protein atoms | 24.0 | 21.2 |
| Water molecules | 33.8 | 35.5 |
| $Zn^{2+}$ | 17.8 | 14.2 |
| Glycerol atoms | 31.6 | 34.4 |

Values in parentheses are statistics for the highest resolution shell

[a] $R_{sym} = \frac{\sum|I - \bar{I}|}{\sum I}$ , with $I$ representing the observed intensity and $\bar{I}$ representing the average intensities for multiple measurements.

[b] $R_{work} = \frac{\sum_{hkl}|F_{obs} - F_{calc}|}{\sum_{hkl}|F_{obs}|}$

[c] $R_{free}$ was calculated as $R_{work}$ but on 5% of the data excluded from the refinement.

[d] calculated by *PROCHECK*[36]

[e] calculated by *MOLEMAN*[37]
